# Inferring seagrass meadow resilience from self-organized spatial patterns

**DOI:** 10.64898/2026.03.24.713893

**Authors:** Àlex Giménez-Romero, Elena del Campo, Manuel A. Matías

## Abstract

Assessing ecosystem resilience at large spatial scales remains a major challenge in ecology and conservation. While resilience is typically inferred from temporal dynamics or perturbation experiments, self-organized ecosystems may encode resilience-related information directly in spatial structure. Here, we combine a mechanistic model of *Posidonia oceanica* self-organization with convolutional neural networks trained on synthetic seascapes to infer meadow condition from single cartographic snapshots. The networks classify model-defined pattern states and estimate an effective mortality parameter along a deterioration gradient. Applied to habitat cartography across the Balearic Islands, the framework revealed regional variation in meadow condition. Cover-preserving randomization altered both classifications and inferred effective mortality, demonstrating dependence on spatial organization beyond vegetation cover alone. Networks trained exclusively on synthetic data transferred to real meadows without site-specific calibration. Our results establish a transferable strategy for integrating ecological theory and machine learning to infer resilience-related ecosystem condition when temporal observations are sparse or unavailable.

## Introduction

Spatial self-organization theory posits that ecosystems governed by scale-dependent competition and facilitation undergo predictable structural shifts under stress, transitioning from homogeneous states through distinct patterned configurations before eventual collapse [1]. Because the geometry of these patterns reflects the system’s underlying ecological condition [2, 3], they have been proposed as critical indicators of ecosystem resilience. For decades, this framework has fueled theoretical research into early warning signals, such as vegetation patchiness and changes in pattern organization [4, 5]. However, despite this robust conceptual foundation, translating these spatial signatures into operational monitoring tools remains a significant challenge.

This challenge exemplifies a broader difficulty in ecology and conservation. Ecosystem resilience— the capacity to absorb environmental stress before shifting to a degraded state—is central to understanding and managing ecological responses to global change [6], yet it remains difficult to quantify at large spatial extents. Most empirical approaches rely on long temporal records [7, 8], observed recovery following perturbations [9], or detailed demographic measurements [10, 11], which are seldom available at policy-relevant scales [12, 13]. Early warning signals of impending ecological transitions—often characterized by critical slowing down and manifesting as rising variance or spatial autocorrelation—offer significant theoretical promise [14, 15]. However, their practical application remains hindered by the need for high-resolution data, and their reliability across ecosystems remains debated [16, 17]. These challenges are especially acute in marine habitat-forming ecosystems, where degradation and recovery unfold over decades [18] and spatially extensive observations are often sparse, fragmented, or temporally inconsistent [19, 20].

In the seagrass *Posidonia oceanica*, mechanistic models developed specifically for this species have shown that scale-dependent feedbacks arising from short-range facilitation and long-range competition can generate large-scale spatial configurations—holes, stripes or labyrinthine structures, and spotted patterns—that emerge and reorganize as mortality increases [21, 22]. Although alternative mechanisms, including inhibitory interactions and auto-toxic self-regulation, can generate related spatial organization [23], the resulting geometric transitions parallel those documented in dryland vegetation under resource stress [3, 24, 25], where spatial patterning has been most extensively theorized as an indicator of resilience and proximity to collapse. Recent comparative work further suggests that these ideas generalize across systems and scales: fairy-circle-like vegetation patterns have now been shown to occur across drylands worldwide, and their presence has been associated with greater temporal stability of vegetation productivity, reinforcing the broader view that self-organized spatial structure can encode functionally meaningful information about ecosystem condition and robustness [26, 27]. More recently, inverse modeling of dryland vegetation patterns has shown that static spatial structure can also be used to infer model parameters and estimate distance to desertification [28]. This perspective is especially relevant because warming-driven stress has been associated with increasing fragmentation of *P. oceanica* meadows at large spatial scales [29], suggesting that spatial organization may encode information about meadow deterioration and resilience-related condition.

Turning that insight into a practical monitoring framework requires bridging two capabilities that have developed largely independently. On the one hand, remote sensing and machine learning are making the large-scale mapping of *P. oceanica* increasingly feasible [30]. On the other hand, existing assessments of meadow condition rely mainly on side-scan sonar surveys complemented by in situ validation, which provide high-quality cartography but are costly and difficult to repeat frequently across broad areas [31, 32]. Improved mapping capacity does not by itself resolve the central ecological challenge: current approaches can map meadow extent and spatial configuration but do not provide direct estimates of ecosystem condition or resilience-related state. Because *P. oceanica* meadows degrade over long timescales, the temporal span of the available high-quality imagery is insufficient to reconstruct deterioration trajectories directly. A critical gap therefore remains between observing seagrass spatial patterns and translating those patterns into actionable information about ecosystem condition.

Here, we address this gap by showing that self-organized spatial patterns can be used to infer seagrass meadow condition from a single cartographic snapshot. Using a previously established mechanistic model of *P. oceanica* self-organization [33], which reproduces the characteristic spatial patterns observed in real meadows and builds on ecological mechanisms with empirical support [21, 23], we generated thousands of synthetic seascapes spanning a deterioration gradient: from continuous meadow cover through progressively fragmented states to meadow collapse. We then trained convolutional neural networks to classify spatial patterns into five discrete states and to estimate the model control parameter *ω* associated with each spatial configuration. This parameter represents the balance between growth and mortality and is hereafter called *effective mortality*. Applying these models to sonar-based habitat cartography from the Balearic Islands, we show that regional-scale variation in meadow condition can be recovered from spatial patterns alone. Although we do not measure resilience in the strict dynamical sense, the inferred discrete states and estimated control parameter together provide a mechanistically grounded proxy for resilience-related ecosystem condition. More broadly, our results show how ecological theory and machine learning can be integrated to monitor ecosystem state in systems where direct observation of deterioration is spatially limited or temporally out of reach.

## Results

### Pattern state and effective mortality are encoded in self-organized spatial structure

Using synthetic seascapes generated by a mechanistic model of *Posidonia oceanica* self-organization, we trained convolutional neural networks to infer both discrete pattern states and the continuous effective mortality parameter from single spatial snapshots (Fig. 1). The model describes the large-scale dynamics of *P. oceanica* meadows, in which clonal growth and spatial interactions generate characteristic spatial patterns as mortality increases (Fig. 1a). Along this mortality gradient, the system shifts from a continuous meadow cover to patterned configurations—meadows with holes, striped or labyrinthine patterns, and seagrass spots—before ultimately collapsing into a no seagrass state. These spatial reorganizations are accompanied by an overall decline in seagrass cover, such that the pattern type and cover jointly reflect the meadow’s position along a mechanistically defined deterioration pathway. Because we apply the model to presence/absence habitat data, we binarized the synthetic seascapes before training (see Methods).

**Figure 1:**
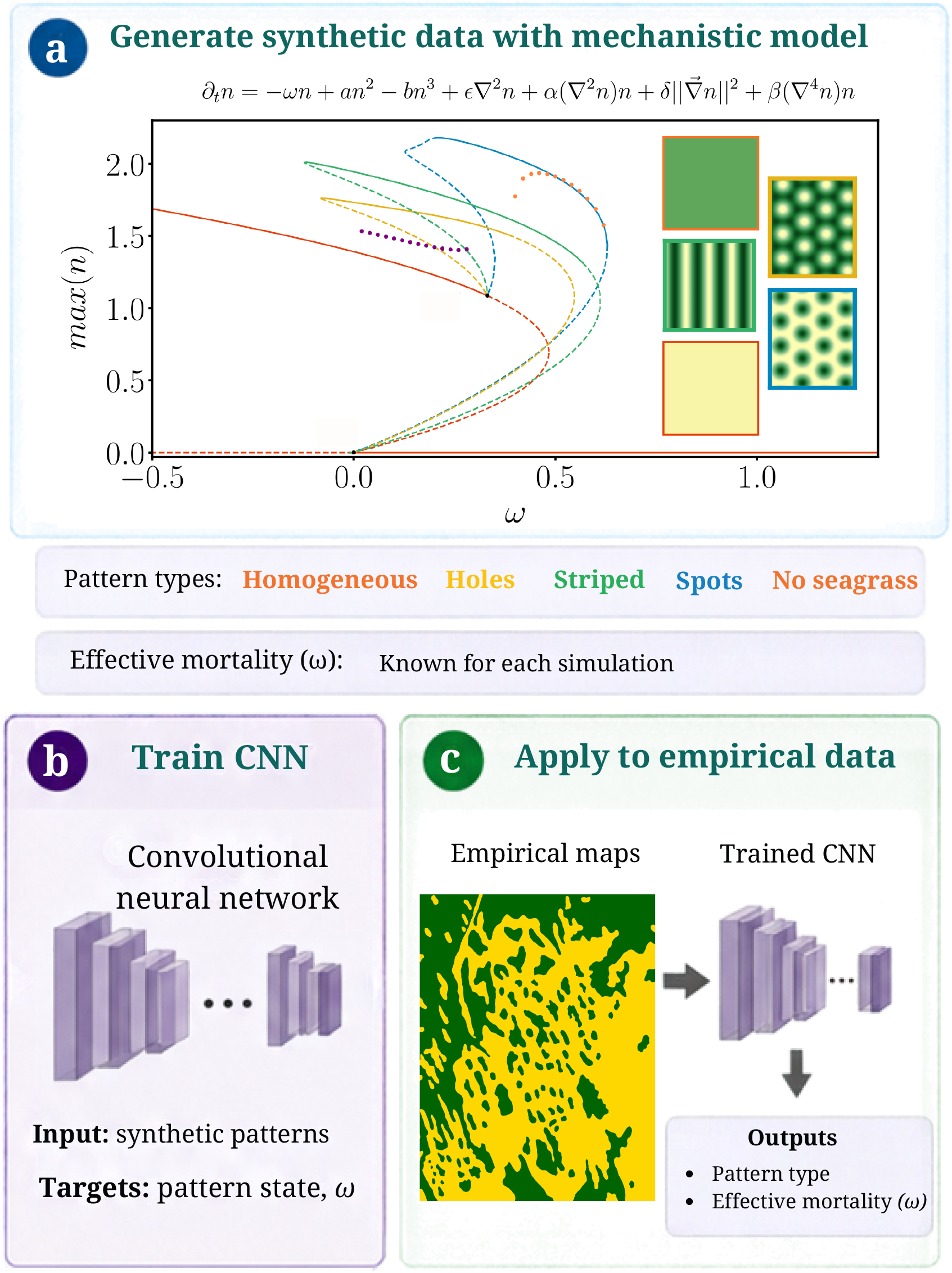
Theory-guided framework for inferring seagrass meadow condition from spatial patterns. (a) Synthetic training data are generated using the mechanistic model of *P. oceanica* self-organization. The bifurcation diagram shows the stationary solutions of the model as effective mortality *ω* increases, including homogeneous meadows and patterned states with holes, stripes or labyrinthine configurations, and spots that precede meadow collapse (reproduced from [33]). Solid and dashed lines denote stable and unstable solution branches, respectively, whereas dotted solutions correspond to localized states, which are not used in this work. Each simulated spatial configuration is therefore associated with both a discrete pattern type and a known value of effective mortality *ω*. (b) Convolutional neural networks are trained exclusively on these synthetic spatial patterns to learn the inverse mapping from spatial structure to the corresponding pattern state and effective mortality. (c) The trained networks are subsequently applied to empirical binary habitat cartography, providing each spatial snapshot with an inferred pattern type and an inferred effective mortality *ω*.

Using synthetic seascapes generated along this gradient, we trained convolutional neural networks to infer meadow condition in two complementary ways (Fig. 1b). First, a classification model assigned each spatial snapshot to one of the discrete pattern states predicted by the theory. Second, a regression model estimated the effective mortality parameter of the mechanistic model underlying each simulated pattern, providing a continuous measure of deterioration. These two outputs are related but not redundant. Each patterned state spans a finite range of effective mortality values, so spatial configurations assigned to the same discrete state can nevertheless correspond to different values of *ω*. The classifier therefore identifies the broad morphological state of the meadow, while the regression resolves its position within that state along the model-defined deterioration pathway. The trained CNN models were subsequently applied to empirical data of the Balearic Islands (Fig. 1c). The classification model recovered the theoretical sequence of spatial states with very high accuracy on the held-out synthetic test set (Supplementary Fig. 11, left). Misclassifications were rare and occurred primarily between neighboring states (i.e., spots and stripes) along the expected deterioration pathway, rather than between structurally distant configurations. This is ecologically important because it indicates that the learned representation preserves the ordered progression from a continuous meadow to increasingly fragmented states. Thus, the spatial motifs generated by the theoretical model are sufficiently distinctive to allow a robust reconstruction of discrete meadow states from spatial patterns alone.

The regression model showed that the same snapshots also encode continuous information about deterioration. The predicted and true effective mortality parameters aligned closely across most of the parameter range (Supplementary Fig. 11, right), showing that self-organized spatial structure can be used to recover a latent ecological parameter in addition to a categorical state label. The largest deviations occurred at low mortality, where binarization removed density variation in the synthetic data, making it impossible to distinguish between homogeneous meadows with different mortality levels. This limitation does not apply to fragmented patterns, where the structural complexity of the spatial arrangement retains enough discriminatory information to differentiate between mortality levels. Therefore, this performance reduction reflects a genuine loss of information in weakly deteriorated patterns rather than a breakdown of the inference framework.

Furthermore, prediction uncertainty differed markedly among spatial states. Inference was most precise for striped meadows, with a 95th-percentile absolute error of 0.034, and remained comparatively precise for meadows with holes and seagrass spots, with corresponding errors of 0.066 and 0.072. Uncertainty was substantially larger for homogeneous meadows (0.193), because *ω* is encoded there through shoot density rather than spatial geometry, and the presence/absence cartography does not record density. Near-homogeneous meadows with different *ω* therefore map onto nearly identical binary configurations, a limitation intrinsic to the data rather than to the binarization step or to class sample size. Empirical mortality estimates are therefore most reliable within the patterned regime and should be interpreted more cautiously for homogeneous meadows (Supplementary Fig. 17 and Table 9).

### Theory-trained models transfer to empirical seagrass cartography

Next, we tested whether the relationship between spatial patterns and meadow condition learned from synthetic seascapes remains detectable in real habitat cartography. This is a stringent test of the framework because empirical maps contain heterogeneity not explicitly represented in the theoretical model, including local environmental variability, cartographic noise, and departures from idealized, stationary patterning. Nevertheless, applying the classifier to empirical maps of the coasts of the Balearic Islands recovered spatial mosaics that were strongly consistent with the expected organization along a deterioration gradient (Fig. 2).

**Figure 2:**
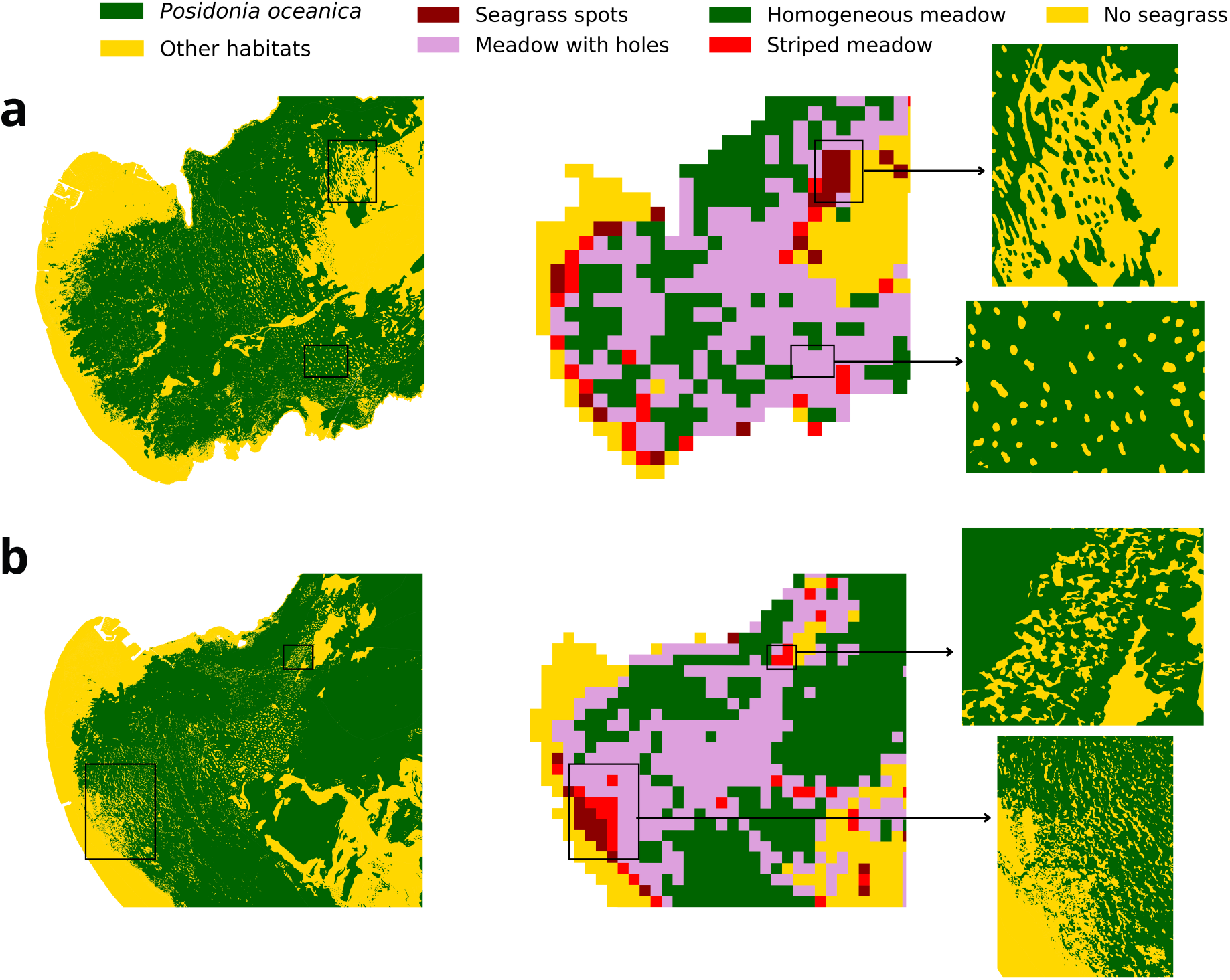
Application of the pattern classifier to empirical habitat cartography. Examples from the coasts of (a) Pollença and (b) Alcúdia in northeast Mallorca. For each site, the left panel shows the empirical habitat map, the central panel shows the inferred spatial state of the meadow, and the right panels show close-up images of representative regions. The classifier recovers heterogeneous mosaics of meadow conditions, including transitions among homogeneous meadows, meadows with holes, striped or labyrinthine states, spotted patterns, and no seagrass, demonstrating transfer from synthetic training data to real-world cartographic patterns.

We classified continuous meadow areas as homogeneous, while assigning more degraded regions patterned states based on their geometry and degree of fragmentation. Meadows containing internal gaps within an otherwise continuous cover were classified as meadows with holes, elongated and fragmented structures as striped or labyrinthine states, and isolated patches as seagrass spots. Close-up views make this transfer particularly clear (Fig. 2). They show that the classifier recovers the same types of spatial signatures used to define the synthetic training set, even though real meadows exhibit greater irregularity than model-generated patterns. This agreement is ecologically important because it indicates that the patterned states predicted by the self-organization model are not merely theoretical abstractions but also capture structural motifs that are observable in empirical seascapes.

The regression model provides a complementary layer of inference by producing spatially explicit maps of the inferred mortality parameter *ω* for the same empirical cartography (Fig. 3; see also Supplementary Figs. 12 to 15). These maps extend the discrete classification into a continuous estimate of deterioration across the seascape, revealing variations in meadow condition among patterned regions. Taken together, the pattern classifications and mortality estimates show that empirical cartography can be translated not only into discrete meadow states but also into a continuous, mechanistically grounded indicator of the ecological state.

**Figure 3:**
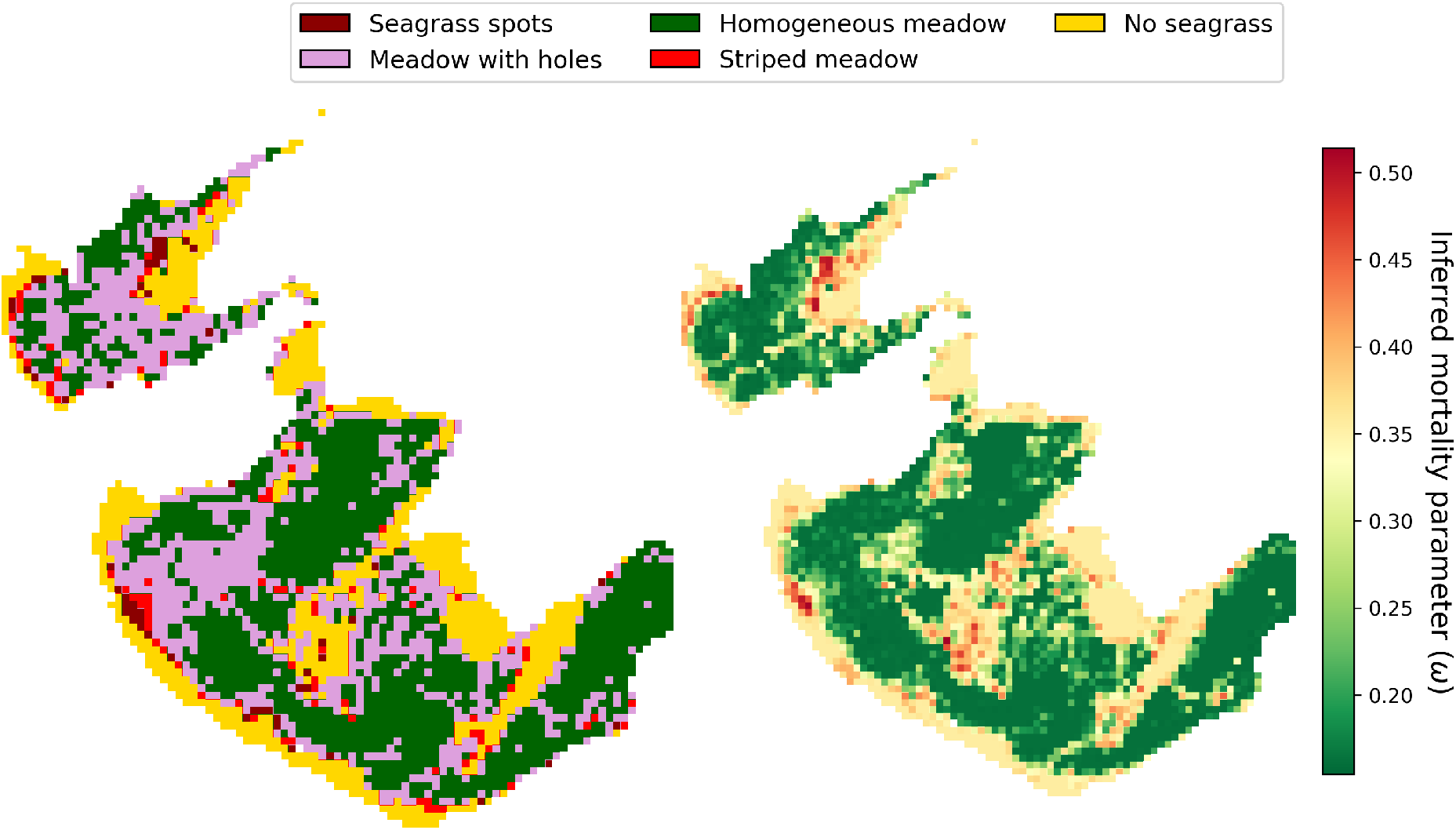
Spatially explicit meadow-state classification and inferred mortality parameter in Pollença and Alcúdia bays in northeast Mallorca. Left: Discrete meadow states inferred by the classification model, including homogeneous meadow, meadow with holes, striped or labyrinthine meadow, seagrass spots, and no seagrass. Right: Corresponding spatially explicit map of the inferred mortality parameter *ω* obtained with the regression model for the same area. The mortality map extends the discrete classification into a continuous estimate of deterioration, revealing fine-scale variation in meadow condition among patterned regions.

This transferability has two implications. First, it shows that synthetic training data derived from mechanistic theory can support ecological inference in real systems. Second, it indicates that the theoretical model captures spatial statistics realistic enough to bridge the gap between self-organization theory and operational monitoring. More broadly, these results show that habitat cartography can be used not only to map where seagrass occurs but also to infer how meadow condition varies across space (Figs. 2 and 3; see also Supplementary Figs. 12 to 15).

### The inferred mortality parameter *ω* captures structural signatures of meadow deterioration

To test whether the inferred states and mortality values corresponded to ecologically meaningful variation in meadow condition, we compared them with complementary structural descriptors derived directly from empirical cartography (Fig. 4). Across all analyses, the framework’s inferred outputs were closely associated with the expected deterioration signatures, indicating that the inferred gradient is consistently reflected in complementary aspects of meadow spatial structure rather than simply in the assigned pattern labels. This interpretation is further supported by the distribution of inferred mortality across degraded pattern classes, which increases from meadows with holes to striped meadows and seagrass spots, consistent with the expected deterioration sequence (Supplementary Fig. 16).

**Figure 4:**
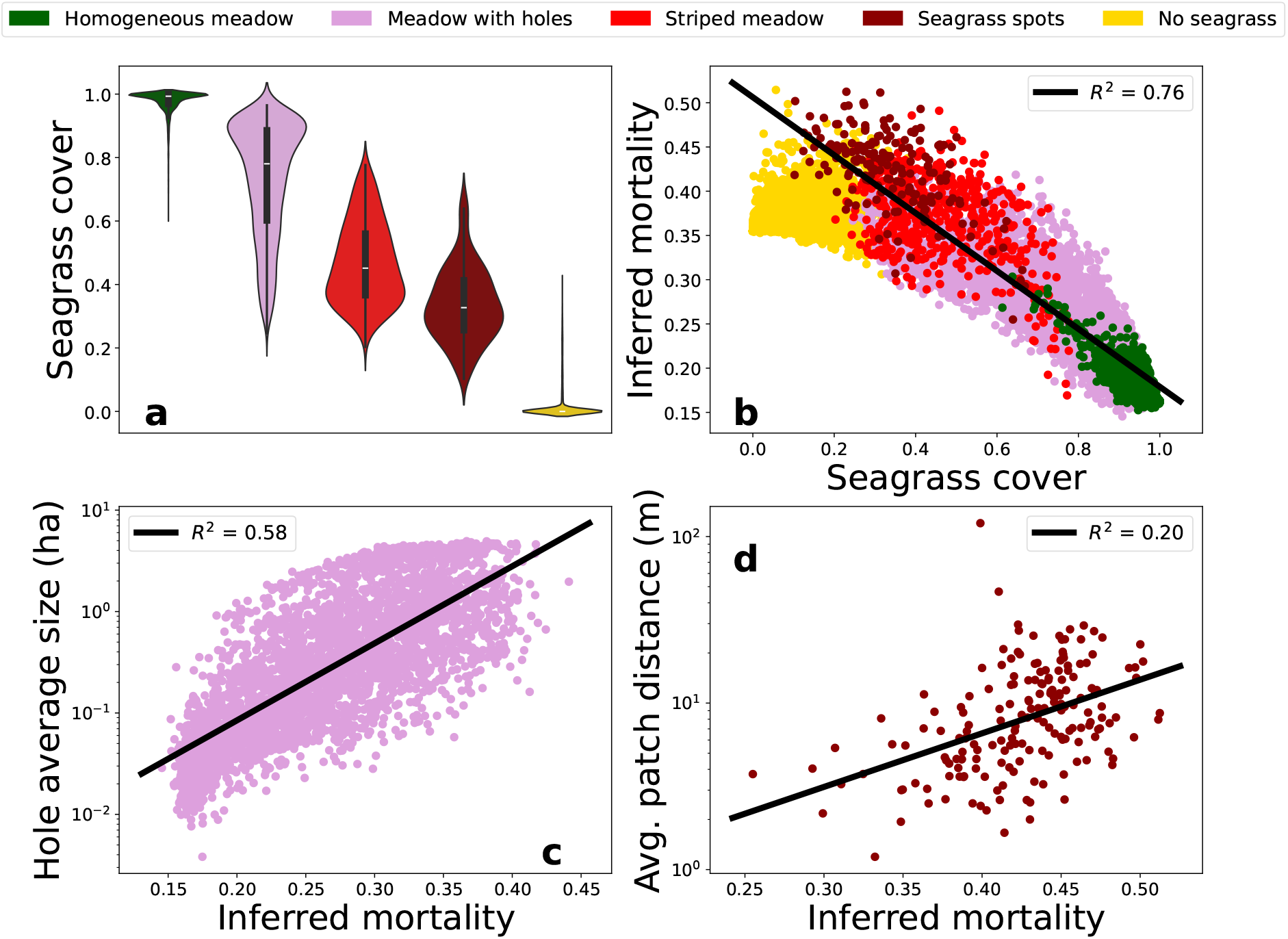
The inferred mortality parameter *ω* captures structural signatures of meadow deterioration. (a) Distribution of seagrass cover across the five inferred spatial states. Cover declines systematically from homogeneous meadows to no seagrass states, although overlap among intermediate states reflects coexistence within the patterned regime. (b) Inferred mortality parameter versus seagrass cover for all mapped samples, showing a strong negative relationship (*R*^2^ = 0.76). (c) Mean hole area versus inferred mortality for samples classified as meadows with holes, indicating that internal gaps enlarge as deterioration increases (*R*^2^ = 0.58). (d) Mean inter-patch distance versus inferred mortality for spotted patterns, showing a weaker but positive association (*R*^2^ = 0.20). Together, these relationships indicate that the inferred mortality is ecologically interpretable and tracks multiple dimensions of meadow structure.

Seagrass cover declined systematically across the inferred spatial states (Fig. 4a). Homogeneous meadows showed the highest cover, followed by meadows with holes, striped or labyrinthine patterns, and spotted states, whereas no seagrass states contained little or no vegetation. Although intermediate-patterned classes showed partial overlap—an expected outcome given that certain parameter combinations allow distinct spatial patterns to coexist (Fig. 1a), [21, 33])—their overall ordering was consistent with the mortality-driven progression predicted by the theoretical model. This result is ecologically important because it shows that the discrete states recovered by the classifier correspond to a broad structural gradient from continuous to increasingly degraded meadow configurations.

The continuous estimate of the mortality parameter captured this gradient more clearly. Across all mapped samples, seagrass cover declined strongly with increasing inferred mortality (Fig. 4b), yielding a marked negative relationship (*R*^2^ = 0.76) consistent with the theoretical model [33]. Thus, the regression-derived mortality parameter is a mechanistically interpretable descriptor of meadow structure that tracks the major axis of ecological degradation. Within the patterned regime, inferred mortality also captured more specific signatures of fragmentation. In samples classified as meadows with holes, the mean hole area increased with the mortality parameter (Fig. 4c), indicating that internal gaps expand as deterioration progresses. Similarly, in spotted meadows, the mean inter-patch distance increased with inferred mortality (Fig. 4d), suggesting that seagrass patches become more isolated as meadow structure breaks down. This is consistent with the increased prevalence of localized structures near the edge of stability in the spotted meadow branch (see Methods).

These within-class relationships illustrate the additional information provided by the continuous inference. For example, two meadows can both be classified as “meadows with holes” while differing in hole size and inferred effective mortality, corresponding to different positions along the same model branch. Likewise, spotted meadows with different degrees of patch isolation can share the same categorical state while differing in inferred *ω*. Effective mortality therefore refines, rather than duplicates, the discrete classification. Because *ω* drives the sequence of spatial states in the mechanistic model, its inferred value also locates a configuration in model-parameter space relative to transitions between spatial states and the limits of the corresponding pattern branch. In addition, these relationships show that the effective mortality parameter *ω* captures multiple complementary dimensions of meadow deterioration, from overall vegetation loss to the size and spacing of the fragmented structures. More broadly, they support interpreting inferred mortality as a continuous, ecologically grounded indicator of meadow condition.

### Inference depends on spatial organization beyond seagrass cover

Because seagrass cover declines along the theoretical deterioration gradient, we tested whether CNN predictions depended on cover alone or also on spatial organization. For each empirical image, we generated a randomized counterpart by reshuffling its pixels, thereby preserving seagrass cover exactly while destroying spatial structure. Applying the same models to these randomized images produced substantial changes in both pattern classification and inferred effective mortality (Fig. 5).

**Figure 5:**
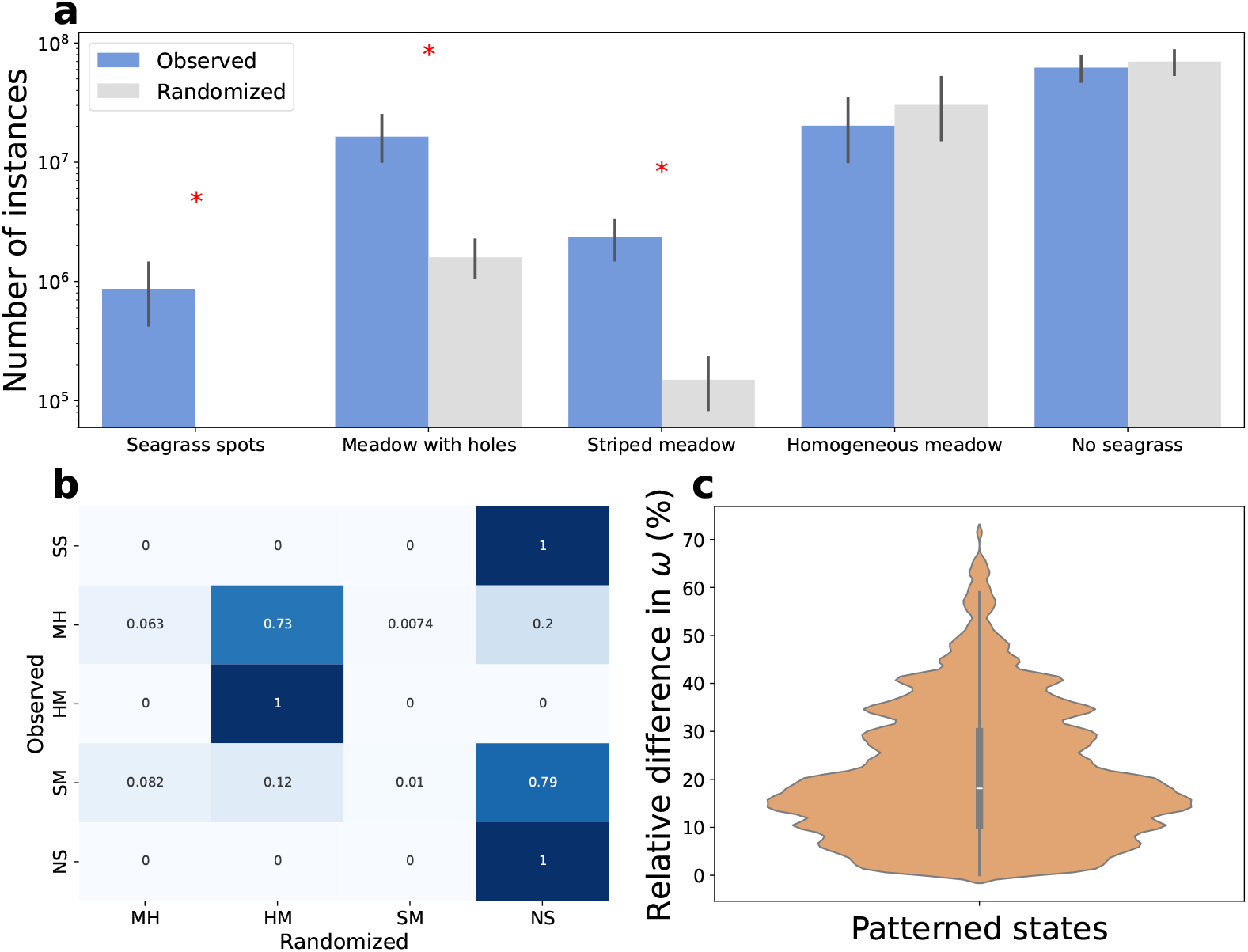
Cover-preserving null-model test of the CNN inference. (a) Mean number of empirical image instances assigned to each spatial state before and after pixel randomization across the study zones; error bars indicate one standard deviation. Randomization preserves seagrass cover exactly while destroying spatial organization. All classes differed significantly between observed and randomized images according to two-sided paired Wilcoxon signed-rank tests with Holm correction (*p ≤* 0.0039). Asterisks highlight the patterned states, where changes were largest. (b) Transition matrix between the class assigned to each observed image and that assigned after randomization. Values indicate the fraction of images within each observed class reassigned to each randomized class. SS: seagrass spots; MH: meadow with holes; HM: homogeneous meadow; SM: striped meadow; NS: no seagrass. (c) Distribution of the relative change in inferred *ω* between observed and randomized images for the patterned states. The marked changes produced by cover-preserving randomization show that both pattern classification and mortality inference depend on spatial organization, not cover alone.

The effect of randomization depended strongly on the initial spatial state. Seagrass-spot, meadow-with-holes, and striped configurations were systematically reassigned once their spatial organization was destroyed, whereas images classified as homogeneous meadow or no seagrass were largely retained. This is expected because the patterned states are defined by non-random spatial organization, including patch aggregation, holes, stripes, and characteristic spatial scales, whereas homogeneous and no-seagrass states contain comparatively little spatial structure beyond their high or low occupancy. Accordingly, state abundance changed most for the patterned classes. Across the 14 study zones, the abundance of all five predicted states nevertheless differed between observed and randomized images (*p ≤* 0.0039; Supplementary Table 8).

Randomization also produced substantial changes in inferred *ω* within patterned meadows (Fig. 5c). Because each observed–randomized pair has identical seagrass cover, vegetation occupancy alone cannot explain these changes. Together, the classification and regression results show that both inference tasks depend on spatial organization beyond fractional cover.

### Regional-scale application reveals spatial variation in meadow condition across the Balearic Islands

Having established that self-organized spatial patterns can be translated into ecologically inter-pretable measures of meadow condition, we applied the framework to the full cartographic dataset of the Balearic Islands to quantify regional variation in the state of *P. oceanica* meadows (Fig. 6). At this scale, the inferred spatial states show that meadow conditions are not uniformly distributed across the archipelago but form a heterogeneous mosaic of continuous, patterned, and degraded configurations.

**Figure 6:**
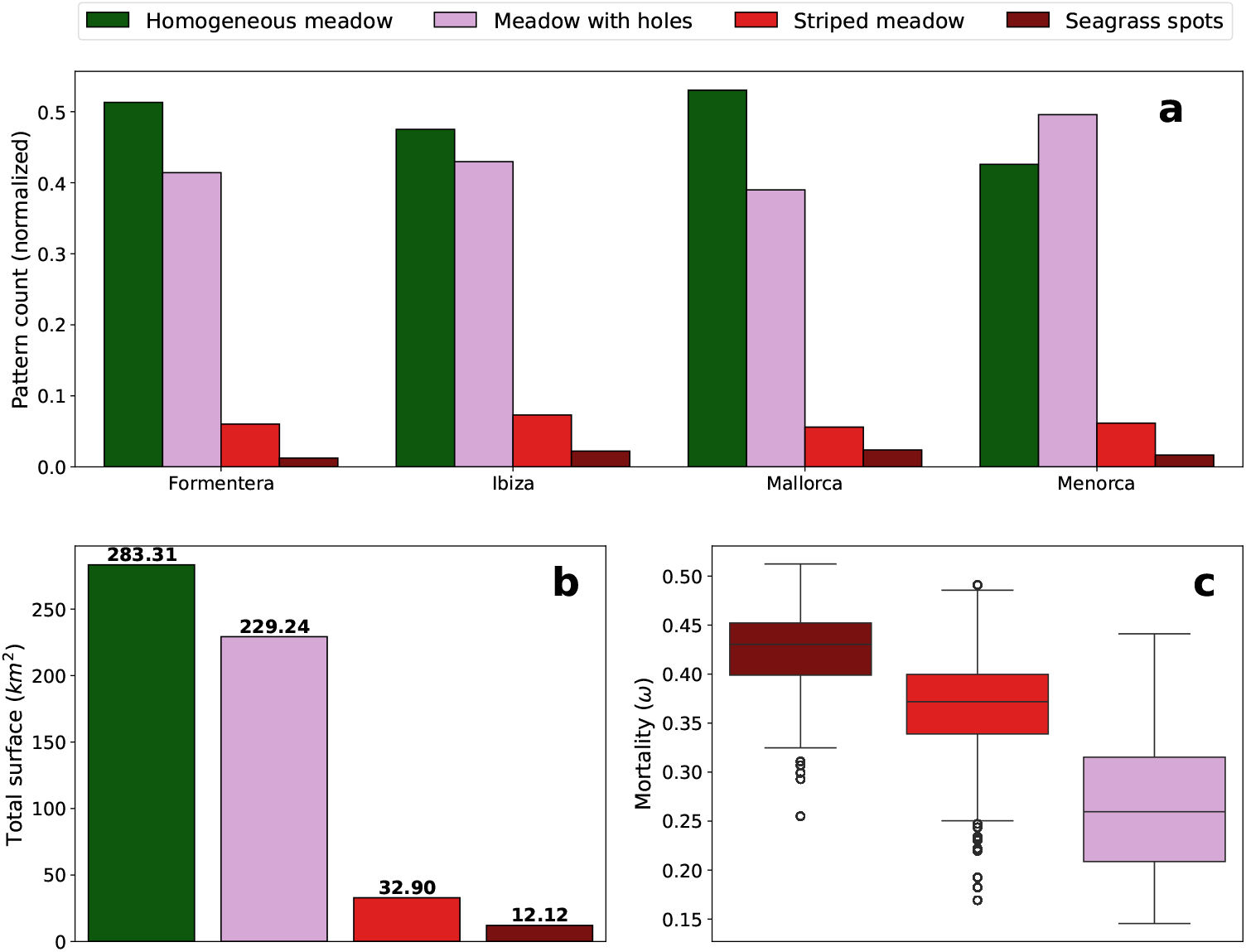
Regional patterns of inferred meadow condition across the Balearic Islands. (a) Relative abundance of the inferred spatial states on each island. (b) Total mapped area occupied by each state across the archipelago. (c) Distributions of inferred mortality parameter *ω* for degraded pattern classes. Homogeneous meadows and meadows with holes dominate the mapped area, but their relative prevalence differs among islands; Menorca shows the largest contribution of meadows with holes, indicating a greater prevalence of an intermediate state along the model-defined deterioration sequence. Across degraded classes, inferred mortality follows the expected ordering, with fragmented meadows showing the highest values.

Homogeneous meadows and meadows with holes were the dominant inferred states across the Balearic Islands, accounting for most of the mapped seagrass area (Fig. 6a,b). However, their relative prevalence varied markedly by island. Homogeneous meadows are generally the most abundant class, except on Menorca, where meadows with holes contribute more. More fragmented states, including striped patterns and seagrass spots, occupied a smaller fraction of the total mapped area but were sufficiently widespread to reveal substantial heterogeneity in meadow conditions at the regional scale. Continuous estimates of mortality *ω* were consistent with the inferred spatial patterns. Across the degraded pattern classes, inferred mortality increased from meadows with holes to striped or labyrinthine states and seagrass spots (Fig. 6c), following the order expected from the mortality-driven sequence predicted by the theoretical model.

## Discussion

Our results show that self-organized spatial patterns carry sufficient information to infer the condition of *Posidonia oceanica* meadows from a single cartographic snapshot. By combining a mechanistic model of seagrass self-organization with deep learning, we show that both the discrete meadow state and an effective mortality parameter can be inferred from spatial structure alone. The inferred effective mortality should be interpreted as a model-based coordinate along the deterioration pathway, rather than as a direct estimate of the temporal rate of meadow decline. This approach therefore provides resilience-related information from static habitat cartography when long-term observations and direct demographic measurements are unavailable, while remaining complementary to repeated surveys required to quantify temporal rates of change. More broadly, the framework addresses a central challenge in ecosystem monitoring: characterizing resilience-related ecosystem condition across broad spatial scales when long-term observations and direct demographic measurements are unavailable.

The key advance of this framework is that model-based information on meadow condition can be extracted from static spatial structure. For slow-growing foundation species such as *P. oceanica*, year-to-year changes may be subtle, making spatial information a potentially valuable complement to conventional temporal monitoring. In the theoretical model, increasing effective mortality reorganizes spatial structure along a characteristic sequence—from continuous cover to holes, striped or labyrinthine states, isolated spots, and ultimately collapse [21, 33]—and we show that this model-defined trajectory is sufficiently imprinted in spatial patterns to be recovered from a single snapshot. Habitat cartography can therefore provide a spatially explicit diagnostic of meadow condition, while repeated observations remain necessary to determine how rapidly that condition is changing through time.

The empirical application across the Balearic Islands reveals a heterogeneous mosaic of continuous, patterned, and fragmented meadows that scalar cover metrics alone cannot capture. The prevalence of hole patterns around Menorca is particularly notable, as this configuration corresponds to an intermediate stage in the theoretical deterioration sequence. However, in practice, such spatial signatures may also arise from differences in geomorphology, substrate properties, coastline complexity, or historical stress regimes, rather than reflecting a more advanced deterioration trajectory per se. The observed contrast therefore points to systematic structural differences among islands, but the ecological drivers remain unresolved. More broadly, the analysis shows that habitat cartography, interpreted through the lens of self-organization theory, can reveal large-scale spatial variation in meadow structure and highlight regions whose patterning departs from the rest of the system.

A central result is that models trained exclusively on synthetic seascapes—without exposure to empirical habitat images—transfer successfully to real maps, recovering coherent spatial states and effective-mortality estimates across the Balearic cartography. This transfer indicates that the mechanistic model generates structural signatures that are also expressed in real meadows, despite the absence of site-specific tuning. We therefore interpret the model as a suitable generative basis for inference, consistent with the independent empirical support for the underlying ecological mechanisms established in previous work [21, 23]. More broadly, the framework strengthens the connection between spatial self-organization theory and operational ecosystem monitoring [16, 34, 35], consistent with recent work showing that spatial patterns can confront mechanistic predictions with empirical data in other self-organized ecosystems [26–28].

The inferred effective mortality further supports this interpretation: its associations with seagrass cover, hole area, and inter-patch distance indicate that the model recovers a continuous, mechanistically grounded gradient of meadow condition rather than discrete visual classes. Importantly, the cover-preserving randomization shows that this inference is not reducible to vegetation cover alone. When spatial organization was destroyed while keeping cover unchanged, both pattern classification and inferred effective mortality changed substantially, particularly for patterned states (Fig. 5). This shows that the CNNs exploit information in the spatial arrangement of vegetation beyond its fractional occupancy. This continuous indicator captures within-class variation and may reveal early shifts toward fragmentation that precede visible structural transitions, although its interpretation as a direct precursor of collapse should be treated cautiously. The state-dependent prediction errors further indicate that inferred effective mortality is most informative within the patterned regime, whereas estimates for homogeneous meadows remain less identifiable from binary cartography alone.

The discrete spatial state and inferred effective mortality should therefore be interpreted as complementary descriptions of meadow condition. The classification identifies the broad morphology of the meadow—for example, whether it contains holes, forms stripes, or is fragmented into spots— whereas effective mortality provides a continuous coordinate along the modelled deterioration pathway. Importantly, each of these spatial states persists over a finite interval of *ω*. Two meadows can therefore display the same broad pattern while occupying different positions within the corresponding model branch. This distinction is reflected in the empirical data: within meadows classified as having holes, hole size increases with inferred *ω*, while within spotted meadows, interpatch distance also increases with *ω*. Thus, the continuous inference captures variation that is not represented by the categorical state alone. This continuous estimate also differs conceptually from a conventional fragmentation metric. Fragmentation describes an observed property of the spatial configuration, whereas *ω* is a latent parameter of the mechanistic model inferred from that configuration. Because increasing *ω* drives the sequence of spatial reorganizations in the model, its inferred value places an observed configuration along the underlying model-defined stress gradient and relative to transitions between spatial states and branch limits. Effective mortality is therefore expected to covary with fragmentation, since both reflect the same deterioration process in the model, but it is not simply another categorical or geometric measure of fragmentation. Accordingly, absolute values of inferred *ω* should not yet be interpreted as empirically calibrated mortality rates or management thresholds; their current interpretation is comparative and conditional on the mechanistic model. Establishing a direct correspondence with field demographic rates would require independent temporal or demographic observations.

Recent studies have increasingly sought to translate self-organized spatial structure into quantitative assessments of ecosystem condition, combining mechanistic models, spatial indicators, remote sensing, and image-based inference [26, 28, 35, 36]. These approaches range from confronting model predictions with empirical spatial patterns to extracting resilience-related indicators and inferring latent model parameters from static observations. Here, CNNs learn spatial features directly from synthetic configurations generated by an established species-specific model, providing an amortized mapping to both pattern state and effective mortality that can be applied across extensive cartographic datasets. Together, these developments illustrate the growing potential of integrating spatial self-organization theory with modern inference methods to extract model-based information on ecosystem condition from spatial observations.

Several limitations should be acknowledged. Regression performance declines at low mortality because the habitat maps used for inference contain only presence–absence information and therefore do not preserve variation in seagrass density. As a result, the model cannot distinguish between near-homogeneous meadows that may correspond to different mortality parameter values but share the same binary configuration, reflecting inherent information loss introduced by binarization [37]. At the opposite end of the deterioration gradient, the weaker relationship between inferred mortality and inter-patch distance in highly fragmented meadows suggests that spatial configuration is not determined exclusively by seagrass self-organization, but may also be modulated by environmental heterogeneity [23]. In addition, the framework is restricted to stationary self-organized patterns and does not capture potentially relevant non-stationary spatiotemporal regimes, such as defect turbulence, spirals, or traveling structures, which may also contribute to resilience [23, 38]. A further limitation arises from the interaction between incomplete cartographic coverage and the patch-based CNN approach. Unmapped areas (no-data values) reduce the effective sample size and introduce spatially heterogeneous prediction coverage, while fixed-size patches generate boundary effects in regions with truncated spatial context. This is particularly relevant for pattern types that depend on mesoscale spatial structure, such as stripe organization or inter-patch spacing, and more generally reflects an implicit scale selection that may limit the representation of multi-scale processes.

Together, these limitations suggest that future developments should move beyond scalar mortality inference and treat meadow condition as a multidimensional property of spatial organization. This could involve integrating richer geometric descriptors of pattern structure, such as landscape fragmentation metrics [39–41], Minkowski functionals [42], or topological summaries of connectivity and patch arrangement, while also incorporating environmental covariates such as bathymetry, substrate, hydrodynamic exposure [43], or thermal stress [29]. Extending the framework to temporal data when available would provide a more direct link to deterioration dynamics, and improving uncertainty quantification would help identify regions where ecological inference is intrinsically weak. Taken together, these extensions would move the framework toward a more complete representation of meadow condition as an emergent, spatially structured, and partially uncertain ecological property.

Although developed here for *P. oceanica*, the framework is conceptually general. It builds on the premise that self-organized spatial patterns provide an observable projection of latent ecological state, a principle widely explored in dryland ecosystems [28, 36]. Our results extend this perspective to seagrass meadows and suggest a general strategy for combining mechanistic models, synthetic data, and machine learning to infer ecosystem condition when direct empirical labels are scarce. Looking ahead, integration with large-scale remote sensing offers substantial potential. Coupled with emerging satellite-based habitat mapping technologies [30], this framework could enable repeated, spatially extensive assessments of *P. oceanica* condition across the Mediterranean basin. Given projected climate-driven habitat loss [44, 45] and increasing fragmentation [29], extracting resilience-related information from single spatial snapshots may provide useful complementary information for monitoring and management.

## Methods

### Theoretical model for seagrass vegetation patterns

To generate the synthetic seascapes used for model training, we employed the reduced one-component model of clonal seagrass dynamics introduced in [33]. This model is a simplified approximation of the more detailed bi-component integro-differential framework developed in [21], which explicitly accounts for rhizome elongation in different directions, branching, shoot mortality, and spatially nonlocal interactions. Although the original model reproduces the main mechanisms underlying seagrass self-organization, its numerical complexity makes large-scale training-data generation computationally demanding. The reduced model retains the key pattern-forming phenomenology of the full framework while providing a more tractable description of meadow dynamics.

The model describes the time evolution of the isotropic vegetation density *n*(*x, y, t*), which is defined as the number of shoots per unit area at position (*x, y*) and time *t* [33]. It is obtained by approximating the nonlocal interaction term of the original nonlocal model [21], through a moment expansion, which yields the following partial differential equation:

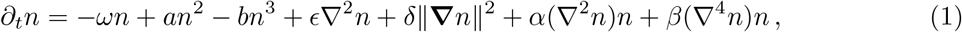

where *∂*_*t*_ denotes the time derivative and ∇ denotes the nabla operator.

This reduced model Eq. (1) provides a generic, mechanistically interpretable description of clonal vegetation dynamics. In particular, it preserves the existence of a bare substrate state, allows for non-negative vegetation density, and reproduces the main sequence of self-organized spatial patterns generated by the full model [21, 33]. Unlike standard vegetation models that are primarily based on seed dispersal, Eq. (1) also includes a term proportional to *∥****∇****∥n* ^2^, which captures the large-scale spatial signature of rhizome-driven clonal growth [21].

The dynamics exhibited by Eq. (1) is controlled by the balance between growth and mortality *ω*, local facilitation *a*, local competition *b*, seed dispersal *ϵ*, clonal growth *δ*, and coefficients *α* and *β* that arise from the first two moments of the nonlocal interaction kernel of [21], as shown in [33]. In the model, *ω* multiplies the linear loss term *™ ωn* and therefore controls the balance between growth and mortality at low vegetation density. We thus refer to this parameter as *effective mortality* throughout. In our study, we treated the effective mortality parameter *ω* as the primary control parameter, as it governs the transition from continuous meadow cover to patterned and ultimately collapsed states. We fixed all remaining parameters to the values reported in [33], as listed in Table 1.

**Table 1.** Parameters of Eq. (1), [46] and their numerical values.

| Parameter | Meaning | Value |
| --- | --- | --- |
| $a$ | local facilitation | 1.3914 |
| $b$ | local competition | 1.0 |
| $\epsilon$ | diffusion | $1.1480 \times 10^{-2}$ |
| $\delta$ | clonal growth | $1.0270 \times 10^{-2}$ |
| $\alpha$ | 1st moment nonlocal interaction term | -1.7668 |
| $\beta$ | 2nd moment nonlocal interaction term | -1.0 |

The scenario exhibited by Eq. (1) is summarized in the bifurcation diagram in Fig. 1(a). A zero solution (bare substrate) is always present and is unstable for *ω <* 0 and stable for *ω >* 0. A homogeneously populated branch is the only stable pattern for negative values of *ω*, where growth dominates, and becomes unstable at the MI point, at which three branches of patterned states emerge, which are initially unstable. These branches are meadows with holes (negative hexagons), yellow line; stripes or labyrinthine patterns, green line; meadows with spots, or patchy meadows (positive hexagons), blue line. These pattern states delay the tipping point at which the meadow disappears at some *ω >* 0 by leading to fragmented states [3]. The panel also shows single dot states, which are localized structures (see also [22]), being relevant the one with a single spot. In this reference, fronts between some of these states appear when they meet. Such fronts can remain static or move, describing the invasion of one biomass configuration by another or its retreat, and can account for the irregular arrangements of spotted patterns near the transition to a barren substrate.

We solved Eq. (1) numerically using a pseudospectral scheme [33], which integrates the linear terms exactly in Fourier space. All simulations were performed on a periodic square domain of size *Lx* = *Ly* = 40 m, discretized on a 256 *×* 256 square grid. Periodic boundary conditions were used, as customary in studies of spatial pattern formation, to characterize the intrinsic self-organized dynamics of the model without introducing effects associated with the finite boundaries of the computational domain. Different boundary conditions may influence the detailed spatiotemporal dynamics near meadow edges, but such boundary-driven effects are outside the scope of the present analysis, which focuses on the characteristic bulk spatial patterns generated by the model. Unless otherwise stated, all parameters except *ω* were fixed to the values reported in Table 1 (with 4-digit precision [46]). These values are consistent with those of the original integro-differential model [21, 33] and were fixed independently of the empirical cartography analysed here. To ensure numerical stability of the nonlinear terms, we applied a standard linear stabilization procedure that leaves the original equation unchanged while improving time-integration behavior (see Supplementary Section 1.2 for details). All simulations were implemented in Julia [47] using FourierFlows.jl [48]. The complete model implementation is openly available in the GitHub and Zenodo repositories [49, 50].

### Pattern dataset generation

We generated the synthetic dataset by numerically integrating Eq. (1) over a range of the mortality parameter *ω*, keeping all other parameters fixed at the values listed in Table 1. Because *ω* controls the transition from continuous vegetation to patterned and ultimately collapsed states, sweeping this parameter provides a mechanistically grounded method for generating spatial snapshots spanning the full deterioration sequence. To trace the branches of patterned solutions efficiently, we used a continuation procedure in which the stationary solution at each value of *ω* served as the initial condition for the next step.

The continuation procedure yielded 259 distinct stationary configurations, including striped (or labyrinthine) and hexagonal states with varying aspect ratios. Data augmentation was applied to increase the dataset size and diversity while preserving the large-scale spatial structure of each pattern and involved translations and noise addition (see Supplementary Section 1.5 for details). Thresholding of the continuous density fields was performed to produce binary representations of vegetation occupancy consistent with empirical presence-absence cartography (see Supplementary Section 1.6 for details).

We then thresholded the simulated density fields into binary presence-absence maps to match empirical habitat cartography and augmented the dataset to increase size and diversity. The final classification dataset comprised 4,591 spatial snapshots across five meadow states: 2,704 striped or labyrinthine patterns, 1,008 filled hexagonal patterns, 432 empty hexagonal patterns, 240 homogeneously vegetated states, and 200 no-seagrass states corresponding to the mechanistic model’s bare-substrate solution. Because the model control parameter *ω* is well-defined only for meadow-bearing states, the 200 no-seagrass images were excluded from the regression task, yielding 4,390 images for effective-mortality inference.

### Deep learning model training and evaluation

We developed two convolutional neural network (CNN) models to infer meadow conditions from spatial patterns. Both models accept a two-dimensional spatial snapshot as input and follow the same convolutional neural-network framework, with architectures optimized separately for classification and regression and task-specific output layers. The first model performs pattern classification, assigning each input image to one of five meadow states: homogeneous meadow, meadow with holes, striped or labyrinthine meadow, seagrass spots, and no seagrass. The second model performs continuous regression, predicting the mortality parameter *ω* associated with each input pattern.

All CNNs were implemented in TensorFlow [51] and trained on a synthetic dataset partitioned into training, validation, and test subsets (70:10:20 split). Although models were trained on both continuous-density and binary presence-absence versions of the dataset, we retained the binary-data models for the main analysis, as they are directly comparable to empirical habitat cartography. Model selection was based on categorical cross-entropy and mean squared error for the classification and regression tasks, respectively; complementary performance metrics and full architectural details are provided in Supplementary Section 2.1.

### Uncertainty in mortality inference

We quantified the expected prediction error of the mortality regressor using the held-out synthetic test set, for which the true value of *ω* is known. Because regression accuracy differs across spatial pattern states, we calculated residuals separately for each pattern class. For each state, we estimated the bias, mean absolute error, root-mean-square error, and the 95th percentile of the absolute prediction error. These quantities provide state-specific uncertainty estimates for empirical predictions. Full methodological details and results are provided in Section 8, Fig. 17, and Table 9.

### Empirical dataset

To apply the trained models to real seascapes, we used the georeferenced benthic habitat cartography of the Balearic Islands provided by the regional government [31, 32]. The benthic cartography covers approximately 2,500 km^2^ of shallow coastal habitat and was produced over roughly two decades through a combination of European and regional monitoring projects, with a major update completed in 2018. It was derived primarily from side-scan sonar, complemented by photo-interpretation of airborne imagery and in situ observations, providing high-resolution representation of benthic spatial configuration. The original dataset comprised 28 habitat classes, which were progressively simplified: first into four broader ecological categories [30, 52], and then, for the present study, into a binary classification distinguishing *Posidonia oceanica* from all other habitat types. This reduction ensures consistency with the binary synthetic dataset used for model training, in which each pixel represents the presence or absence of *Posidonia*. The resulting maps provide spatially explicit information on meadow occupancy and fragmentation but not on shoot density, so they are directly comparable to the thresholded synthetic patterns and serve as the empirical input for inferring meadow state and the mortality parameter *ω* across the Balearic Islands.

### Spatial indicators

To evaluate whether the inferred meadow states and mortality parameter *ω* corresponded to ecologically meaningful variations in seagrass structure, we quantified three spatial indicators directly from the empirical binary maps: seagrass cover, mean hole area, and mean inter-patch distance. These indicators capture complementary aspects of meadow organization, from overall vegetation occupancy to fragmentation geometry. Seagrass cover captures broad meadow deterioration, whereas hole area and inter-patch distance characterize fragmentation geometry within the patterned regime. Full definitions and computational details are provided in Supplementary Section 4.4.

### Cover-preserving spatial randomization

To test whether CNN inference depended solely on seagrass cover, we generated a randomized counterpart of each empirical image by randomly permuting its binary pixels. This procedure preserves image dimensions and seagrass cover exactly while destroying spatial organization, including patch aggregation, holes, stripes, and characteristic spatial scales. We processed observed and randomized images using the same classification and regression models. We evaluated differences in the abundance of predicted states across the 14 study zones using two-sided paired Wilcoxon signed-rank tests, with Holm correction for the five comparisons. Full statistical results are reported in Supplementary Table 8.

## Data availability

The derived data products supporting the findings of this study are available in the Zenodo repository associated with this work [50]. The archive includes region-level geospatial outputs of predicted pattern classes and inferred effective mortality, a pixel-level table of predicted pattern types and mortality-parameter values, and a patch-level dataset containing structural descriptors and inferred quantities.

The original benthic habitat cartography of the Balearic Islands was obtained from the corresponding regional data sources [31, 32]. Access to these raw data is subject to the conditions established by the original data providers. An interactive web application for visualizing the predicted spatial pattern classes is publicly available at [53].

## Code availability

The code used to generate the synthetic patterns, train the deep-learning models, and apply them to empirical habitat cartography is publicly available in the GitHub repository associated with this study [49]. A permanently archived version of the code, together with the trained pattern-classification and effective-mortality regression models, is available through Zenodo [50].

## Author contributions

A.G.R., E.d.C., and M.A.M. conceived the study and contributed to its design. A.G.R. and E.d.C. developed the methodology and performed the analyses. E.d.C. implemented the computational framework and deep-learning models. All authors wrote and revised the manuscript.

## Acknowledgements

This work was supported by grants TED2021-131836B-I00 (SEDIMENT) funded by the Spanish Ministry of Science and Innovation MICIU/AEI/10.13039/501100011033 and by the European Union NextGenerationEU/PRTR Program; PID2021-123723OB-C22 (CYCLE) and PID2024-156062OB-I00 (CHANGE-ME) funded by MICIU/AEI/10.13039/501100011033 and by ERDF, EU; and CEX2021-001164-M (María de Maeztu Program for Units of Excellence in R&D) funded by MICIU/AEI/10.13039/501100011033. A.G.R. acknowledges financial support from grant JDC2024-053275-I, funded by MICIU/AEI/10.13039/501100011033 and FSE+; and from grant PID2021-124731NB-I00, funded by the Spanish Ministry of Science and Innovation MICIU/AEI/10.13039/501100011033 and by ERDF, EU.

We acknowledge Damià Gomila, Pablo Moreno-Spiegelberg, and Daniel Ruiz-Reynés for useful comments and help with the numerical integration of the model [33, 46]. We also acknowledge Nuria Marbà and Elvira Mayol for their valuable comments and suggestions, and Marcial Bardolet and Jorge Moreno from the Species Protection Service of the Government of the Balearic Islands, who provided the habitat data used in this study.

