## Supplementary Information for "Inferring seagrass meadow resilience from self-organized spatial patterns"

**5 Model test results** **19**

**6 Pattern and mortality inference** **20**

**7 Comparison with null model** **23**

**8 State-specific uncertainty of inferred mortality** **24**

[†] These authors contributed equally.

### 1 Pattern generation

#### 1.1 Theoretical model

The theoretical patterns used in this study were generated from the reduced one-component model of clonal seagrass dynamics introduced in [1]. This equation was proposed as a simplified approximation of the more general bi-component integro-differential model developed in [2], which explicitly describes rhizome elongation in different directions, branching, shoot mortality, and spatially nonlocal interactions at the landscape scale. Although the original model captures the main mechanisms underlying seagrass self-organization, its numerical complexity makes large-scale pattern generation computationally demanding. The reduced model retains the essential pattern-forming phenomenology of the full framework while providing a much simpler and more tractable description of vegetation dynamics.

The model describes the time evolution of vegetation density $n(x, y, t)$, which is defined as the number of shoots per unit area at position $(x, y)$ and time $t$. It reads

$$\partial_t n = -\omega n + an^2 - bn^3 + \epsilon \nabla^2 n + \alpha(\nabla^2 n)n + \delta \|\boldsymbol{\nabla} n\|^2 + \beta(\nabla^4 n)n\,, \tag{1.1}$$

where $\partial_t$ denotes the partial derivative with respect to time and $\nabla$ denotes the nabla operator.

This equation is obtained by approximating the nonlocal interaction term of the original model through a moment expansion [1]. The reduced formulation was designed to preserve the main qualitative features of clonal seagrass systems: the existence of a bare-soil solution, nonnegative vegetation densities, and the emergence of spatially patterned states under changing mortality conditions. Importantly, it also incorporates a term proportional to $\|\boldsymbol{\nabla} n\|^2$, which captures the distinctive large-scale signature of rhizome-driven clonal growth [2] and distinguishes this framework from standard vegetation models based primarily on seed dispersal.

The terms in Eq. (1.1) have a direct ecological interpretation. The linear term $-\omega n$ represents a parameter that regulates local linear mortality, representing the net balance between losses and gains at low density. A negative value of $\omega$ indicates net growth. The quadratic and cubic terms $an^2$ and $-bn^3$ describe local facilitative and competitive interactions, respectively. The diffusive term $\epsilon\nabla^2 n$ accounts for local spatial spreading, which can be interpreted as an effective contribution of seed dispersal. The nonlinear gradient term $\delta\|\boldsymbol{\nabla} n\|^2$ captures the contribution of rhizome-driven clonal growth to vegetation expansion. Finally, the terms $\alpha(\nabla^2 n)n$ and $\beta(\nabla^4 n)n$ correspond to the first two moments of the nonlocal interaction kernel and encode the leading effects of spatially extended plant–plant interactions.

Therefore, the dynamics of the system are governed by the mortality parameter $\omega$, local facilitation $a$, local competition $b$, seed dispersal $\epsilon$, clonal growth $\delta$, and the coefficients $\alpha$ and $\beta$ associated with the moment expansion of the nonlocal interaction term. The meaning and numerical values of these parameters are summarized in Supplementary Table 1.

The authors of [2], in which spatial coupling includes a nonlocal integral term, have also published an alternative model [3], endorsed by experimental measurements, in which spatial coupling occurs through self-inhibitory feedback by sulfides generated by the decomposition of dead *Posidonia*. In this case, the observed structures are not stationary patterns but excitable structures. These structures evolve and move at an extremely slow speed of the order of decades (cf. Fig. 3 in [3]) and turn out to be stationary for the purposes of our study.

**Supplementary Table 1.** Definition and characteristics of the parameters of Eq. (1.1).

| **Parameter** | **Definition** | **Characteristics** |
|---|---|---|
| $\omega$ | Linear local mortality | Mortality - growth rates |

| | | |
|---|---|---|
| $a$, $b$ | Local facilitative and competitive interactions parameters | As the equation is written, a positive value of $a$ implies a facilitative interaction (the greater the number of organisms, the greater the benefit to the rest), reducing the death rate at higher density. On the other hand, the constant $b$ (which must be positive for a solution with maximum homogeneous density to exist) represents a competitive interaction with the opposite effect, increasing the death rate at higher densities. |
| $\alpha$, $\beta$ | Interaction terms parameters | Included in order to respect the empty meadow solution ($n = 0$) and always positive density requirements. |
| $\epsilon$ | Diffusive term parameter | Accounts for plant propagation by seed dispersion. |
| $\delta$ | Clonal growth parameter | Term added to previous work. It allows that when $\delta > 0$, the density grows when the gradient is nonzero. This is exclusive to plants that reproduce clonally via rhizome elongation. |

#### 1.2 Details of numerical method to solve the PDE

To solve Eq. (1.1) we consider a square grid with $256 \times 256$ points, and integrate the time evolution of $n(x, y, t)$ using a pseudospectral method, in which the linear terms in Fourier space are integrated exactly, avoiding the artificial dissipation introduced by finite-difference discretizations, while the nonlinear terms are integrated using a second-order in time scheme as described in Ref. [4]. Simulations used a time step, $dt = 10^{-4}$, and typically required on the order of $10^6$ time steps to reach a stationary state, depending on the value of $\omega$ and the resulting pattern type. Striped and labyrinthine patterns are treated as a single class in the analysis, though they differ structurally: true stripes (also called rolls in Pattern Formation Theory [5]) are characterized by a well-defined wavenumber $k$, whereas labyrinthine patterns involve multiple wavenumbers arising through secondary instabilities and become more prevalent at higher values of $\omega$ in the domain of existence of striped patterns. Simulations of labyrinthine patterns can require very long integration times because they exhibit a slow reorganization dynamics.

The numerical stability of the nonlinear terms — particularly $(\nabla^4 n)n$ — was ensured by introducing a linear stabilization term, $+\gamma\nabla^4 n$ in the nonlinear update and a compensating term, $-\gamma\nabla^4 n$ in the linear operator. This procedure leaves the original equation unchanged but improves the numerical behavior of the time integration. The stabilization parameter $\gamma$ was chosen adaptively from the homogeneous vegetated equilibrium of the system. When the non-zero stable homogeneous solution existed, we set

$$\gamma = -\beta n_{\text{stab}},$$

where $n_{\text{stab}}$ denotes the stable homogeneous steady state (Section 1.3 of this SI). For parameter values for which this state did not exist, that is, when $\omega > a^2/(4b)$, we used $\gamma = -2\beta$, which provided stable numerical integration across the explored parameter range. Parameter values are reported to four-digit precision [6], unless those reported in [1].

#### 1.3 Stable homogeneous solution

The homogeneous stationary solutions of Eq. (1.1) correspond to states that are uniform in space and constant in time. They are obtained by setting the temporal and spatial derivatives in Eq. (1.1) to zero, which yields

$$0 = -\omega n + an^2 - bn^3 = -n\left(\omega - an + bn^2\right). \tag{1.2}$$

One solution is the bare state,

$$n = 0 \ . \tag{1.3}$$

The non-zero homogeneous states are obtained by solving

$$\omega - an + bn^2 = 0, \tag{1.4}$$

which is a quadratic equation in $n$. Its two roots are

$$n_{\pm} = \frac{a \pm \sqrt{a^2 - 4b\omega}}{2b} = \frac{a}{2b} \pm \sqrt{\left(\frac{a}{2b}\right)^2 - \frac{\omega}{b}} \, . \tag{1.5}$$

These solutions exist provided that the discriminant is non-negative, that is,

$$a^2 - 4b\omega \geq 0, \tag{1.6}$$

or equivalently,

$$\omega \leq \frac{a^2}{4b} \ . \tag{1.7}$$

Thus, non-zero homogeneous equilibria exist only below the threshold $\omega = a^2/(4b)$.

In the parameter regime considered here, the relevant vegetated homogeneous solution is the larger root,

$$n_{\rm stab} = \frac{a}{2b} + \sqrt{\left(\frac{a}{2b}\right)^2 - \frac{\omega}{b}} \tag{1.8}$$

which corresponds to the stable non-zero branch of the homogeneous dynamics. This quantity is used to define the stabilization parameter $\gamma$ in the numerical scheme described in the Methods section.

For $\omega > a^2/(4b)$, the non-zero homogeneous solution does not exist. In this regime, $n_{\rm stab}$ was replaced by the constant value 2 when estimating $\gamma$ and constructing random initial conditions, as this choice provided stable numerical performance in practice.

#### 1.4 Mortality parameter sweeps

To generate the initial conditions used for the mortality sweeps, we started with random perturbations around the stable homogeneous vegetated solution $n_{\rm stab}$. Specifically, for each grid point we set

$$n_0(x, y) = n_{\rm stab} + 0.1 \times \xi(x, y), \tag{1.9}$$

where $\xi(x, y)$ is a normally distributed random number. Random values were generated in Julia using the default implementation of the Xoshiro256++ algorithm [7]. An example of such an initial condition, together with the corresponding frequency distribution of density values, is shown in Supplementary Fig. 1.

For striped patterns, different spatial configurations can arise for the same value of $\omega$, depending on the initial condition. In particular, horizontal stripes, vertical stripes, more complex structures such as concentric patterns, and labyrinthine structures may all appear in the striped-pattern branch in [1]. Pure stripes, vertical and horizontal, exhibit a single wavenumber $k$, whereas in labyrinths, several $k$'s are relevant, and all of them are stripe-like. To capture this variability, we first generated 150 patterns at $\omega = 0.42$ using random initial conditions, as described above. Four examples are shown in Supplementary Fig. 2. For each simulation, $1.2 \times 10^6$ time steps were performed. Fourteen of the resulting patterns were discarded as defective.

A second striped pattern dataset was generated by sweeping $\omega$ across a range of values. Starting from the upper-left striped pattern shown in Supplementary Fig. 2, we performed sweeps in increments of 0.01, both upward from $\omega = 0.42$ to $\omega = 0.44$ and downward from $\omega = 0.42$ to $\omega = 0.12$. For each value of $\omega$, the system was evolved for $10^6$ time steps before saving the pattern and updating $\omega$. In all cases, the initial condition for each new step of the sweep was the stationary pattern obtained at the previous value of $\omega$. This process yielded 33 additional striped patterns. Labyrinthine structures are striped patterns that exhibit secondary instabilities

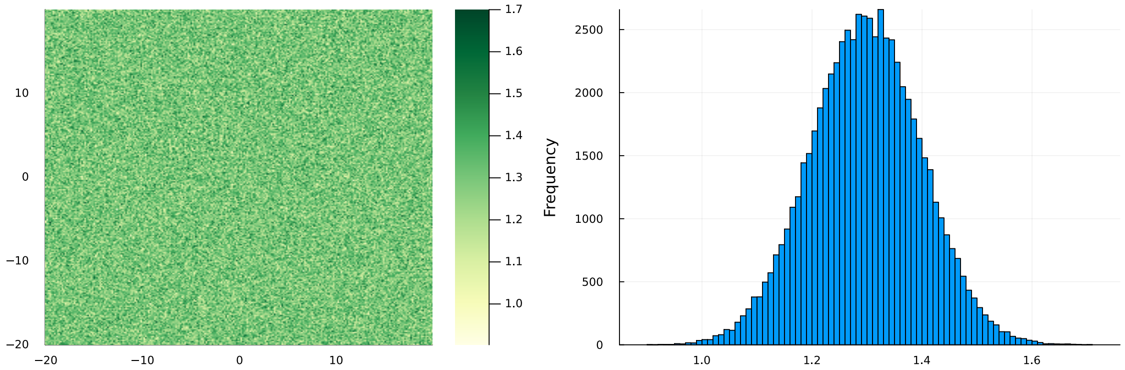

**Supplementary Fig. 1.** Example of the random initial condition used to obtain a homogeneous vegetated meadow for $\omega = 0.12$, with $n_{\text{stab}} = 1.2975$. The accompanying histogram shows the distribution of vegetation density values across the grid.

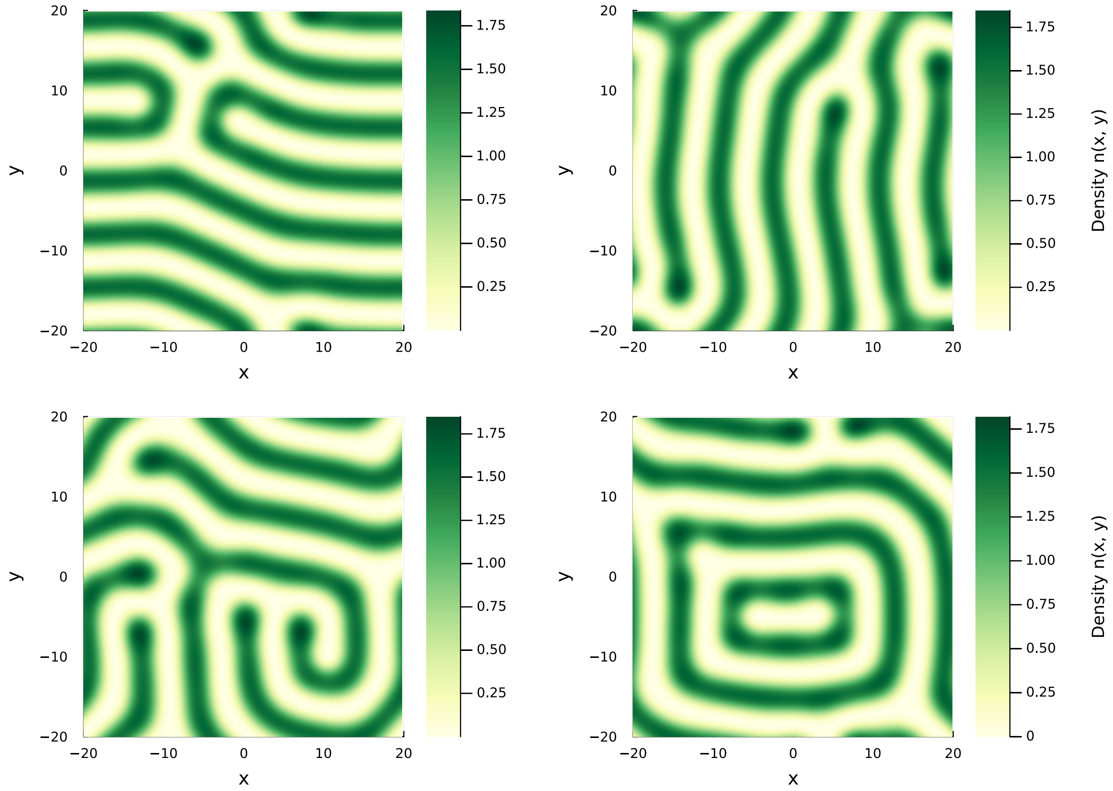

**Supplementary Fig. 2.** Examples of striped (partly labyrinthine) meadow patterns obtained for $\omega = 0.42$. Different spatial organizations can emerge at the same mortality value depending on the initial condition.

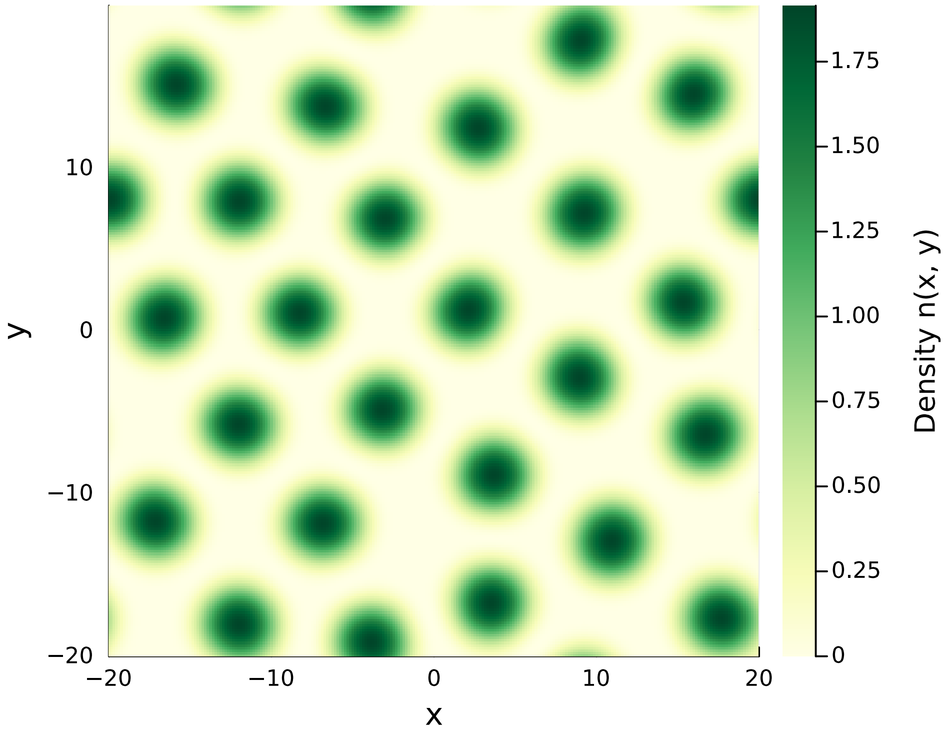

**Supplementary Fig. 3.** Example of a seagrass spot pattern obtained at $\omega = 0.50$, used as the starting point for the corresponding mortality sweep.

with several participating wavenumbers $k$. They exhibit slow convergence, and may not be fully stationary after an established number of time steps.

For the seagrass spot patterns, we first generated a single pattern at $\omega = 0.5$ using a random initial condition constructed as described above. This pattern, shown in Supplementary Fig. 3, was then used as the starting point for a mortality sweep. Specifically, we decreased $\omega$ from 0.5 to 0.24 in steps of $-0.005$ and increased it from 0.5 to 0.55 in steps of 0.005. For each value of $\omega$, the system was evolved for $2 \times 10^6$ time steps before saving the pattern and changing $\omega$. As in the striped case, each new simulation used the pattern obtained in the previous sweep step as an initial condition. This procedure produced 63 spotted seagrass patterns.

For the meadow with hole patterns, we first generated an initial condition by inverting one of the seagrass spot patterns and allowing it to stabilize into a meadow-with-holes pattern at $\omega = 0.2$. The resulting pattern is shown in Supplementary Fig. 4. Starting from this configuration, we performed a sweep in $\omega$ using increments of 0.005 from $\omega = 0.2$ to $\omega = 0.235$ and decrements of $-0.005$ from $\omega = 0.2$ to $\omega = 0.105$. For each value of $\omega$, the model was integrated for $2 \times 10^6$ time steps before saving the pattern and updating the mortality parameter.

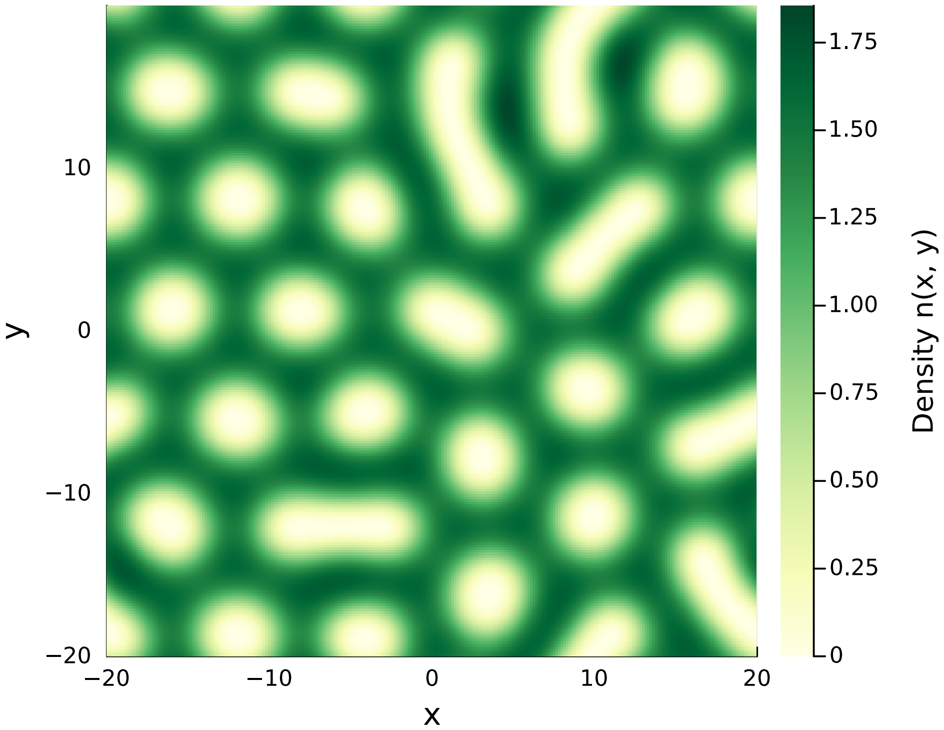

**Supplementary Fig. 4.** Example of a meadow-with-holes pattern obtained at $\omega = 0.20$, used as the starting point for the corresponding mortality sweep.

##### 1.5 Data augmentation

The number of available synthetic patterns differed substantially among the pattern types. In particular, striped patterns were the most abundant, as they display a wide range of geometries and therefore provided the largest diversity of training examples. In contrast, the homogeneous and empty-meadow states showed much lower internal variability and were represented by fewer simulations. This imbalance was accounted for during model training to avoid biasing the neural networks toward the most abundant classes.

For all fragmented pattern classes (striped, seagrass spots, and meadow with hole patterns), we applied the same data augmentation strategy; examples are shown in Supplementary Figs. 5 to 7. In each case, the upper-left panel corresponds to one of the original patterns obtained directly from the numerical simulations. From each original pattern, we generated augmented samples by applying random translations in the $x$ and $y$ directions and by adding noise via random swaps of pixel pairs in the density matrix. The number of swaps was chosen randomly between 1 and the total number of pixels in the image. This procedure increased the size of the dataset by a factor of 16.

An important feature of this augmentation scheme is that the amount of added noise was not fixed but varied from image to image. Consequently, some augmented samples remained similar to the original pattern, whereas others became noticeably noisier. This variability enriched the training set by exposing the models to a broader range of realizations within each pattern class.

For the homogeneous vegetated state, the stable homogeneous solution given by Eq. (1.8) was used to generate the base images. Forty initial images were generated by evaluating this solution for mortality values $\omega$ ranging from 0 to 0.4 in increments of 0.01. For each value of $\omega$, five additional images were created by adding noise, yielding a total of 240 images. This noise was introduced by selecting a random number of pixels between 1 and approximately one-third of the total number of pixels and assigning them random values between 0 and the corresponding homogeneous density. An example of this procedure is shown in Supplementary Fig. 8.

For the empty-meadow state, we followed an analogous procedure. First, 50 images were

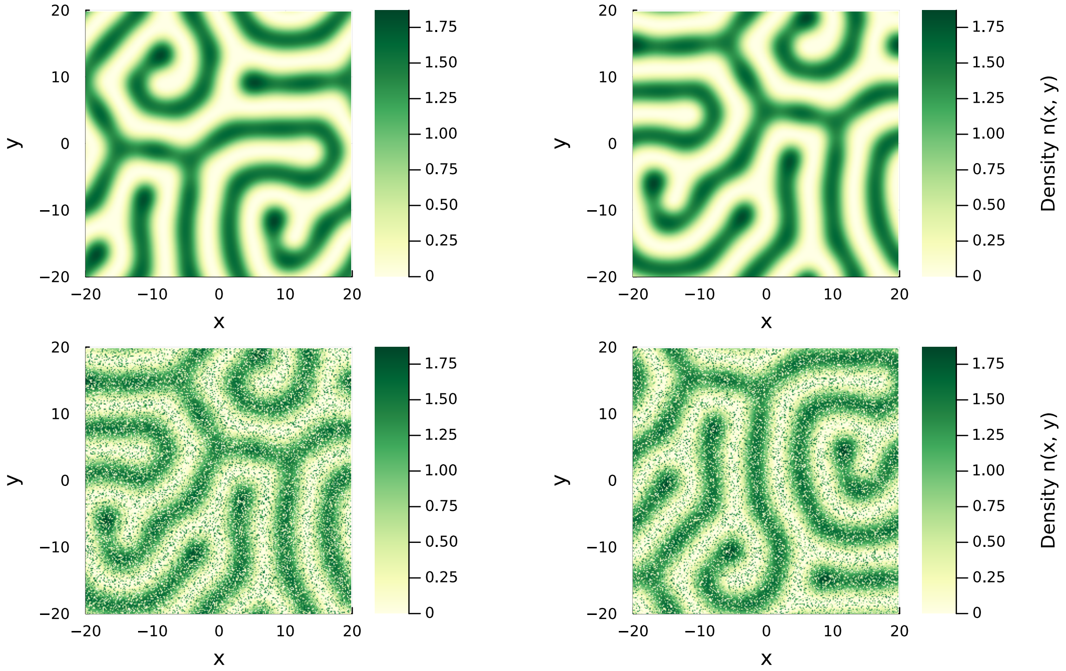

**Supplementary Fig. 5.** Example of the data augmentation procedure applied to a labyrinthine (striped) pattern. The upper-left panel shows the original simulated pattern, and the remaining panels show augmented versions obtained through random spatial translations and pixel swaps.

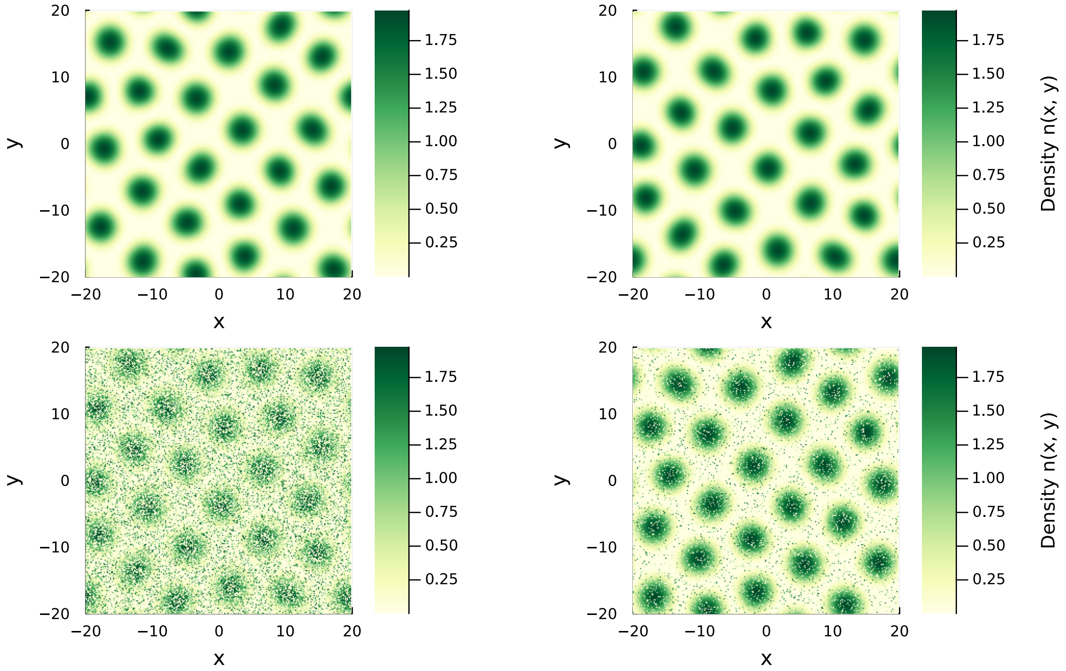

**Supplementary Fig. 6.** Example of the data augmentation procedure applied to a seagrass-spot pattern. The upper-left panel shows the original simulated pattern, and the remaining panels show augmented versions obtained through random spatial translations and pixel swaps.

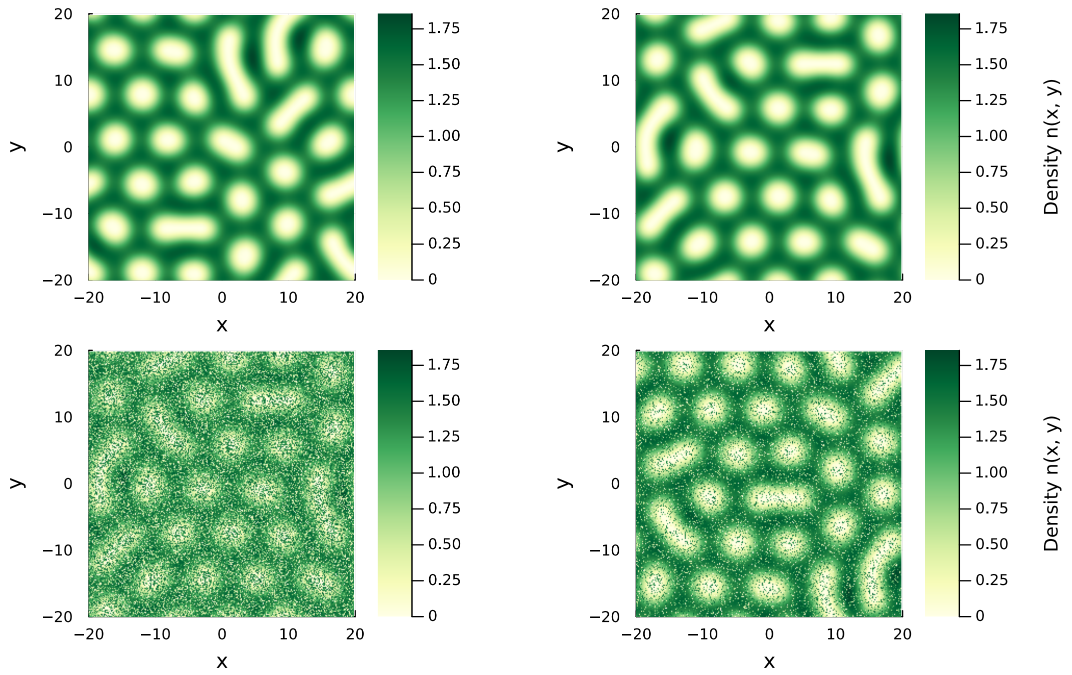

**Supplementary Fig. 7.** Example of the data augmentation procedure applied to a meadow-with-holes pattern. The upper-left panel shows the original simulated pattern, and the remaining panels show augmented versions obtained through random spatial translations and pixel swaps.

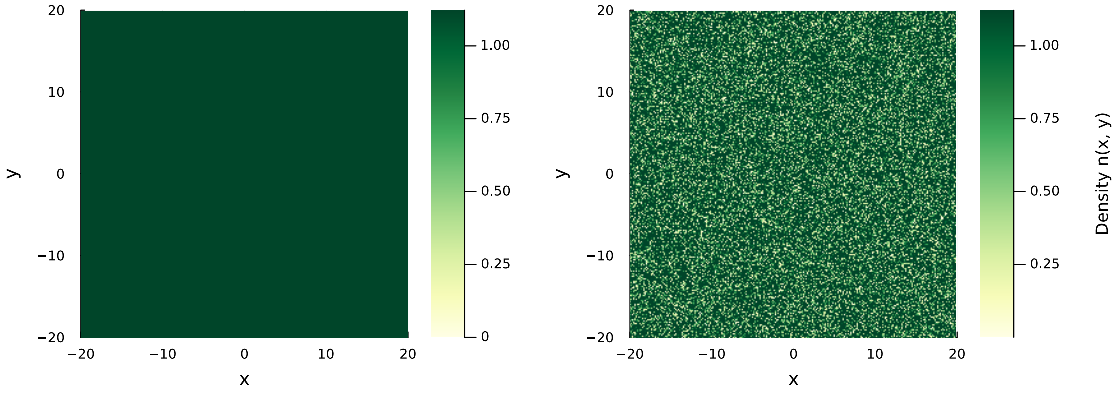

**Supplementary Fig. 8.** Example of the data augmentation procedure applied to a homogeneous vegetated pattern. Starting from the homogeneous solution, additional samples were generated by perturbing a random subset of pixels.

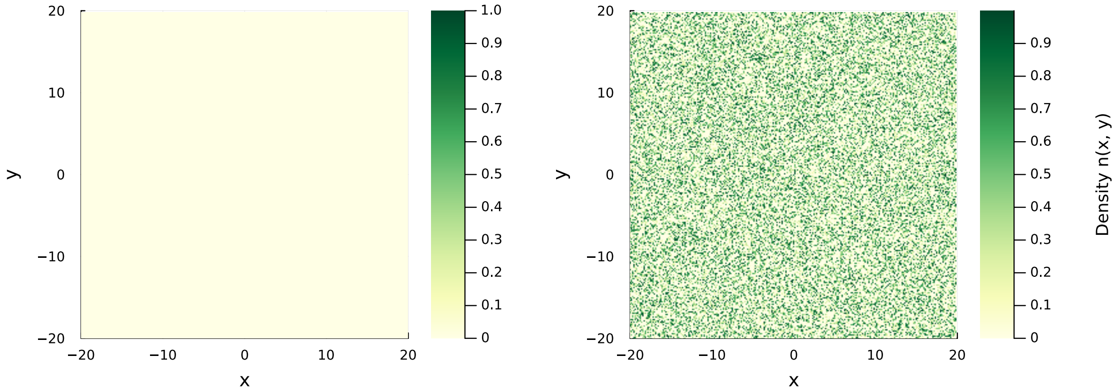

**Supplementary Fig. 9.** Example of the data augmentation procedure applied to an empty-meadow pattern. Starting from the zero-density state, additional samples were generated by perturbing a random subset of pixels.

generated, with all pixels set to zero density. We then generated 150 additional images by adding noise, resulting in 200 samples. Noise was introduced by selecting a random number of pixels between 1 and approximately one-third of the total number of pixels and assigning random values between 0 and 1 to them. An example is shown in Supplementary Fig. 9.

#### 1.6 Thresholding

The empirical habitat cartography used in this study provides information on the spatial distribution of *P. oceanica* as benthic habitat classes but does not include large-scale maps of shoot density. To make the model-generated patterns comparable with these empirical data, we processed the model-generated dataset to represent only the presence or absence of seagrass.

To this end, the continuous vegetation density fields generated by the mechanistic model were thresholded using a density value of 0.4. This value was selected because simulated patterns exhibit a clear separation between vegetated regions, where shoot densities are substantially higher, and unvegetated regions, where densities typically remain below 0.1. The threshold therefore provides a convenient distinction between vegetation presence and absence, although its precise value is not expected to be critical within this range of density separation. Pixels with densities above 0.4 were assigned a value of 1 (presence), whereas those below this threshold were assigned a value of 0 (absence). This procedure was applied only to the synthetic patterns, as the empirical habitat cartography already provides categorical information distinguishing *P. oceanica* from other habitats.

Empty-meadow patterns did not require thresholding because they already correspond to the zero-density state and, therefore, did not depend on the mortality parameter. After thresholding, the binary dataset contained 4,390 patterns. Four representative examples of the resulting binary patterns are shown in Supplementary Fig. 10.

Throughout this Supplementary Information, we refer to the thresholded dataset as the *binary dataset* and to the original density-based dataset as the *continuous dataset*.

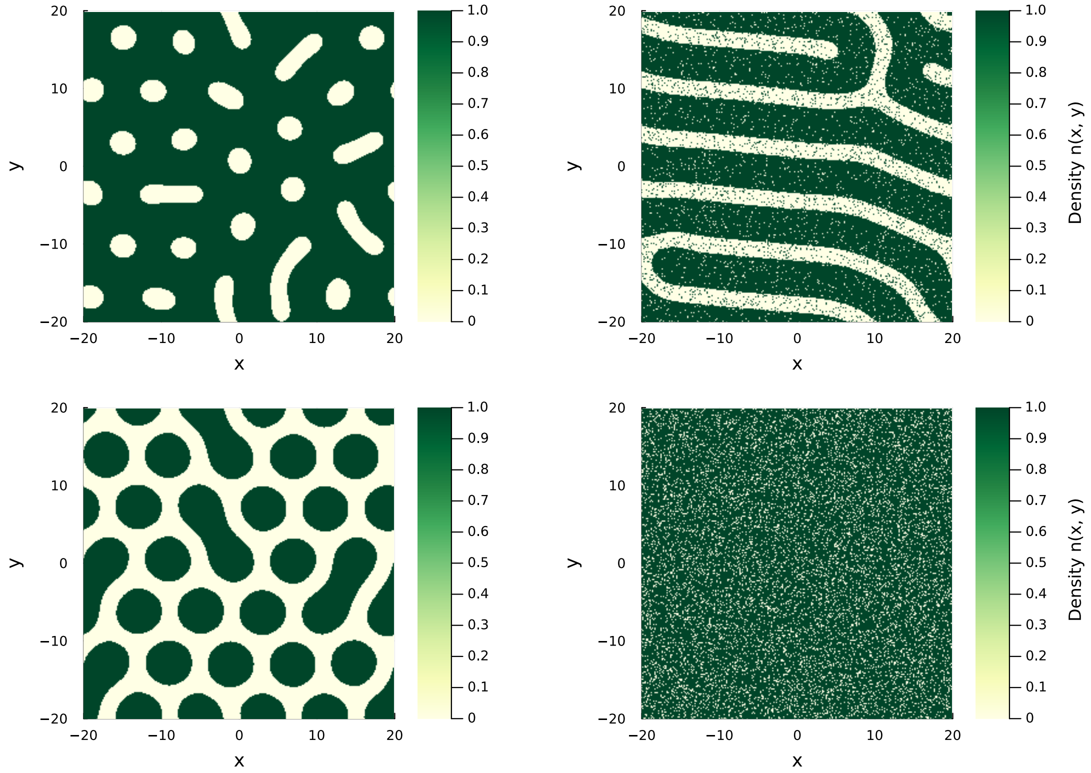

**Supplementary Fig. 10.** Examples of binary patterns obtained after thresholding the continuous density fields. Shown are representative snapshots of a meadow with holes (top left), a striped pattern (top right), a seagrass-spot pattern (bottom left), and an empty meadow (bottom right).

### 2 Deep learning models

The main objective of this study is to infer the spatial state of *Posidonia oceanica* meadows from habitat maps, thereby formulating the problem as an image analysis task. To this end, we employed convolutional neural networks (CNNs) [8], a class of deep learning models widely used for image classification and pattern recognition tasks because of their ability to learn hierarchical spatial features directly from pixel data [9, 10]. Starting from the input image, a CNN progressively extracts increasingly complex features ranging from local edges and textures to higher-level spatial structures and uses them to perform the final prediction.

To identify suitable model configurations, we explored a range of architectures and training hyperparameters. In particular, we varied the number of convolutional layers from 2 to 4, the number of units in the final fully connected layer from 32 to 64, the batch size from 16 to 32, and the number of steps per epoch from 10 to 20. In total, we trained and evaluated 24 models for the pattern-classification task on binary data and 24 on continuous data. For the mortality-estimation task, we trained and evaluated 16 models on binary data and 12 models on continuous data, yielding a total of 76 models.

#### 2.1 Deep learning model training and evaluation procedure

To identify appropriate architectures for each task, we explored candidate CNN configurations through systematic hyperparameter tuning. In total, 48 models were trained for pattern classification and 28 for mortality estimation, varying the number of convolutional layers, the size of the final fully connected layer, and selected training parameters. All models used rectified linear unit (ReLU) activation in hidden layers and were trained for 30 epochs with a learning rate of $10^{-3}$. The training set was used to optimize model parameters, the validation set to select among candidate architectures, and the held-out test set to evaluate final predictive performance. For classification, target labels were encoded as one-hot vectors of length five; complementary performance metrics included mean intersection-over-union and multi-class ROC AUC. For regression, the primary optimization metric was mean squared error; mean absolute error and mean squared logarithmic error were reported as complementary measures. All metrics were computed using `scikit-learn` [11]. Full details of the hyperparameter search and metric definitions are provided in Section 2.2.

**Supplementary Table 2. Composition of the synthetic datasets used for CNN training and evaluation.** Number of samples per spatial pattern state in the training, validation, and held-out test subsets. The no-seagrass state was included in the classification task but excluded from the effective-mortality regression because no unique value of $\omega$ is associated with this state.

| Pattern state | Training | Validation | Test | Total |
|---|---|---|---|---|
| *Pattern classification* | | | | |
| Seagrass spots | 703 | 88 | 217 | 1008 |
| Meadow with holes | 298 | 43 | 91 | 432 |
| Homogeneous meadow | 179 | 18 | 49 | 246 |
| Striped meadow | 1885 | 296 | 523 | 2704 |
| No seagrass | 148 | 14 | 39 | 201 |
| Total | 3213 | 459 | 919 | 4591 |
| *Effective-mortality regression* | | | | |
| Seagrass spots | 703 | 94 | 211 | 1008 |
| Meadow with holes | 287 | 43 | 102 | 432 |
| Homogeneous meadow | 170 | 19 | 57 | 246 |
| Striped meadow | 1913 | 283 | 508 | 2704 |
| Total | 3073 | 439 | 878 | 4390 |

The synthetic datasets were partitioned into training, validation, and held-out test subsets using a 70:10:20 split. Because the number of simulated configurations differed among spatial pattern states, their representation in each subset was not uniform. The exact composition of the datasets used for classification and effective-mortality regression is reported in Supplementary Table 2.

#### 2.2 Hyperparameter tuning

We trained the 76 deep-learning models described in the Methods section and compared their performance on the training and test datasets. The results are summarized in Supplementary Tables 3 to 6, where models are ordered from best to worst according to their performance on the test set, using categorical accuracy (CA) for classification and mean squared error (MSE) for mortality estimation.

Overall, the classification models achieved consistently strong performance. In most cases, the categorical accuracy on the test set was close to or above 90%, indicating that the models reliably distinguished pattern classes, even when evaluated on previously unseen data. Notably, the reduction from continuous density maps to binary presence-absence data did not result in a substantial loss of classification performance.

The model performance generally improved with architectural complexity. In particular, networks with more convolutional layers tended to perform better, although the gains in classification accuracy were often modest; therefore, simpler models with fewer layers could still achieve comparable results. A similar pattern was observed for the mortality estimation models, although the dependence on network depth was more pronounced. In particular, architectures with only two convolutional layers were generally insufficient to achieve good regression performance. Therefore, for the binary dataset, additional models with four convolutional layers were trained to reduce the final prediction error.

**Supplementary Table 3.** Results of categorical accuracy, mean one-hot encoding IoU, and ROC AUC score for 24 pattern classification models with continuous data ordered from highest to lowest categorical accuracy for the test dataset. Their basic parameters have been included: Units of the Last Dense layer (U.L.D.) and Number of Convolution layers (N.C.).

| **Model** | **Parameters** | | **Categ. accuracy** | | **Mean one-hot IoU** | | **ROC AUC** | |
|---|---|---|---|---|---|---|---|---|
| | U.L.D. | N.C. | Train | Test | Train | Test | Train | Test |
| **18** | **32** | **4** | **0.9969** | **0.9956** | **0.9960** | **0.9948** | **0.8642** | **0.8647** |
| 22 | 64 | 4 | 1.0000 | 0.9946 | 1.0000 | 0.9935 | 0.9608 | 0.9589 |
| 3 | 32 | 3 | 0.9906 | 0.9924 | 0.9871 | 0.9767 | 0.9846 | 0.9814 |
| 14 | 64 | 2 | 0.9937 | 0.9859 | 0.9785 | 0.9782 | 0.8740 | 0.8660 |
| 24 | 64 | 4 | 0.9969 | 0.9956 | 0.9959 | 0.9948 | 0.9710 | 0.9689 |
| 8 | 64 | 3 | 0.9812 | 0.9859 | 0.9464 | 0.9735 | 0.9667 | 0.9635 |
| 2 | 32 | 3 | 0.9766 | 0.9837 | 0.8457 | 0.9505 | 0.9682 | 0.9618 |
| 17 | 32 | 4 | 0.9750 | 0.9815 | 0.9592 | 0.9691 | 0.9513 | 0.9474 |
| 5 | 64 | 3 | 0.9812 | 0.9815 | 0.9546 | 0.9644 | 0.9269 | 0.9197 |
| 1 | 32 | 3 | 0.9750 | 0.9750 | 0.9678 | 0.9678 | 0.9144 | 0.9107 |
| 23 | 64 | 4 | 0.9625 | 0.9717 | 0.9159 | 0.9384 | 0.9533 | 0.9489 |
| 21 | 64 | 4 | 0.9812 | 0.9717 | 0.9776 | 0.9681 | 0.9190 | 0.9148 |
| 13 | 64 | 2 | 0.9844 | 0.9652 | 0.9634 | 0.9602 | 0.9033 | 0.8806 |
| 16 | 64 | 2 | 0.9750 | 0.9652 | 0.9698 | 0.9424 | 0.9832 | 0.9710 |
| 20 | 32 | 4 | 0.9812 | 0.9641 | 0.9777 | 0.9531 | 0.9663 | 0.9616 |
| 12 | 32 | 2 | 0.9656 | 0.9619 | 0.9335 | 0.9140 | 0.8821 | 0.8697 |
| 19 | 32 | 4 | 0.9438 | 0.9402 | 0.7391 | 0.7282 | 0.9029 | 0.8994 |
| 7 | 64 | 3 | 0.9438 | 0.9380 | 0.9407 | 0.8456 | 0.9423 | 0.9373 |
| 6 | 64 | 3 | 0.9953 | 0.9303 | 0.9941 | 0.8922 | 0.9261 | 0.9191 |
| 15 | 64 | 2 | 0.9937 | 0.9293 | 0.9795 | 0.8533 | 0.9435 | 0.9352 |
| 11 | 32 | 2 | 0.9125 | 0.9195 | 0.7126 | 0.7449 | 0.8788 | 0.8682 |
| 9 | 32 | 2 | 0.9312 | 0.9119 | 0.7438 | 0.7278 | 0.9453 | 0.9241 |
| 4 | 32 | 3 | 0.9125 | 0.8836 | 0.7288 | 0.6184 | 0.9500 | 0.9440 |
| 10 | 32 | 2 | 0.9922 | 0.8607 | 0.9846 | 0.7135 | 0.9065 | 0.8812 |

**Supplementary Table 4.** Results of Mean Squared Error (MSE), Mean Absolute Error (MAE), and Mean Squared Logarithmic Error (MSLE) for 12 regression models with continuous data ordered from highest to lowest MSE for the test dataset. Their basic parameters have been included: Units of the Last Dense layer (U.L.D.) and Number of Convolution layers (N.C.).

| Model | Parameters | | MSE | | MAE | | MSLE | |
|---|---|---|---|---|---|---|---|---|
| | U.L.D. | N.C. | Train | Test | Train | Test | Train | Test |
| **10** | **32** | **4** | **0.00071** | **0.00093** | **0.0205** | **0.0236** | **0.00044** | **0.00057** |
| 11 | 64 | 4 | 0.00158 | 0.00179 | 0.0319 | 0.0345 | 0.00099 | 0.00110 |
| 12 | 64 | 4 | 0.00217 | 0.00248 | 0.0345 | 0.0375 | 0.00127 | 0.00144 |
| 4 | 64 | 3 | 0.00270 | 0.00307 | 0.0431 | 0.0453 | 0.00153 | 0.00172 |
| 9 | 32 | 4 | 0.00460 | 0.00514 | 0.0495 | 0.0519 | 0.00258 | 0.00285 |
| 1 | 32 | 3 | 0.00372 | 0.00644 | 0.0453 | 0.0614 | 0.00219 | 0.00365 |
| 3 | 64 | 3 | 0.00784 | 0.00850 | 0.0783 | 0.0813 | 0.00452 | 0.00492 |
| 6 | 32 | 2 | 0.00496 | 0.01371 | 0.0500 | 0.0916 | 0.00282 | 0.00776 |
| 7 | 64 | 2 | 0.01170 | 0.01452 | 0.0901 | 0.1009 | 0.00683 | 0.00836 |
| 2 | 32 | 3 | 0.03520 | 0.03371 | 0.1706 | 0.1660 | 0.02046 | 0.01962 |
| 5 | 32 | 2 | 0.12541 | 0.12212 | 0.3373 | 0.3322 | 0.086030 | 0.08400 |
| 8 | 64 | 2 | 0.12628 | 0.12299 | 0.3386 | 0.3335 | 0.08674 | 0.08470 |

**Supplementary Table 5.** Results of categorical accuracy, mean one-hot encoding IoU and ROC AUC score for 24 pattern classification models with binary data ordered from highest to lowest categorical accuracy for the test dataset. Their basic parameters have been included: Units of the Last Dense layer (U.L.D.) and Number of Convolution layers (N.C.).

| Model | Parameters | | Categ. accuracy | | Mean one-hot IoU | | ROC AUC | |
|---|---|---|---|---|---|---|---|---|
| | U.L.D. | N.C. | Train | Test | Train | Test | Train | Test |
| **18** | **32** | **4** | **0.9922** | **0.9880** | **0.9900** | **0.9858** | **0.9327** | **0.9291** |
| 22 | 64 | 4 | 1.0000 | 0.9869 | 1.0000 | 0.9845 | 0.9337 | 0.9282 |
| 24 | 64 | 4 | 0.9969 | 0.9837 | 0.9962 | 0.9807 | 0.9447 | 0.9428 |
| 17 | 32 | 4 | 0.9656 | 0.9837 | 0.9599 | 0.9709 | 0.9706 | 0.9677 |
| 6 | 64 | 3 | 0.9922 | 0.9815 | 0.9787 | 0.9735 | 0.9123 | 0.9069 |
| 20 | 32 | 4 | 0.9937 | 0.9815 | 0.9925 | 0.9686 | 0.8972 | 0.8952 |
| 12 | 32 | 2 | 0.9969 | 0.9804 | 0.9845 | 0.9578 | 0.9433 | 0.9300 |
| 3 | 32 | 3 | 0.9969 | 0.9782 | 0.9766 | 0.9782 | 0.9617 | 0.9564 |
| 19 | 32 | 4 | 0.9875 | 0.9771 | 0.9544 | 0.9735 | 0.9066 | 0.9030 |
| 5 | 64 | 3 | 0.9875 | 0.9771 | 0.9838 | 0.9590 | 0.9341 | 0.9256 |
| 10 | 32 | 2 | 0.9969 | 0.9684 | 0.9961 | 0.9484 | 0.9137 | 0.9024 |
| 9 | 32 | 2 | 0.9937 | 0.9608 | 0.9767 | 0.9348 | 0.9514 | 0.9298 |
| 13 | 64 | 2 | 0.9906 | 0.9608 | 0.9884 | 0.9175 | 0.9510 | 0.9297 |
| 7 | 64 | 3 | 0.9812 | 0.9565 | 0.9778 | 0.9218 | 0.9522 | 0.9451 |
| 8 | 64 | 3 | 0.9625 | 0.9543 | 0.9179 | 0.9503 | 0.9696 | 0.9605 |
| 2 | 32 | 3 | 0.9734 | 0.9478 | 0.9678 | 0.9184 | 0.8773 | 0.8725 |

| | | | | | | | | |
|---|---|---|---|---|---|---|---|---|
| 1 | 32 | 3 | 0.9402 | 0.9402 | 0.9363 | 0.8812 | 0.9255 | 0.9181 |
| 21 | 64 | 4 | 0.9375 | 0.9042 | 0.8791 | 0.8701 | 0.9360 | 0.9334 |
| 14 | 64 | 2 | 1.0000 | 0.8988 | 1.0000 | 0.8561 | 0.9910 | 0.9497 |
| 4 | 32 | 3 | 0.9875 | 0.8934 | 0.9855 | 0.6539 | 0.8503 | 0.8453 |
| 16 | 64 | 2 | 0.9969 | 0.8836 | 0.9962 | 0.8238 | 0.9678 | 0.9330 |
| 23 | 64 | 4 | 0.8750 | 0.8585 | 0.7362 | 0.8479 | 0.9464 | 0.9458 |
| 11 | 32 | 2 | 0.8687 | 0.7573 | 0.6397 | 0.5834 | 0.9286 | 0.8910 |
| 15 | 64 | 2 | 0.8875 | 0.7280 | 0.7171 | 0.5300 | 0.8581 | 0.8162 |

**Supplementary Table 6.** Results of Mean Squared Error (MSE), Mean Absolute Error (MAE), and Mean Squared Logarithmic Error (MSLE) for 12 regression models with binary data ordered from highest to lowest MSE for the test dataset. Their basic parameters have been included: Units of the Last Dense layer (U.L.D.) and Number of Convolution layers (N.C.).

| **Model** | **Parameters** | | **MSE** | | **MAE** | | **MSLE** | |
|---|---|---|---|---|---|---|---|---|
| | U.L.D. | N.C. | Train | Test | Train | Test | Train | Test |
| **14** | **32** | **5** | **0.00117** | **0.00146** | **0.0198** | **0.0232** | **0.00078** | **0.00098** |
| 16 | 64 | 5 | 0.00174 | 0.00208 | 0.0298 | 0.0327 | 0.00106 | 0.00129 |
| 15 | 64 | 5 | 0.00168 | 0.00208 | 0.0289 | 0.0321 | 0.00105 | 0.00132 |
| 10 | 32 | 4 | 0.00273 | 0.00367 | 0.0390 | 0.0474 | 0.00162 | 0.00216 |
| 11 | 64 | 4 | 0.00312 | 0.00379 | 0.0406 | 0.0470 | 0.00191 | 0.00231 |
| 12 | 64 | 4 | 0.00346 | 0.00440 | 0.0397 | 0.0470 | 0.00206 | 0.00260 |
| 13 | 32 | 5 | 0.00413 | 0.00450 | 0.0547 | 0.0571 | 0.00235 | 0.00259 |
| 3 | 64 | 3 | 0.00261 | 0.00483 | 0.0406 | 0.0582 | 0.00155 | 0.00281 |
| 2 | 32 | 3 | 0.00124 | 0.00522 | 0.0233 | 0.0579 | 0.00077 | 0.00301 |
| 4 | 64 | 3 | 0.00197 | 0.00592 | 0.0306 | 0.0618 | 0.00121 | 0.00343 |
| 9 | 32 | 4 | 0.00562 | 0.00627 | 0.0624 | 0.0672 | 0.00319 | 0.00359 |
| 1 | 32 | 3 | 0.03635 | 0.03460 | 0.1737 | 0.1671 | 0.02118 | 0.02020 |
| 7 | 64 | 2 | 0.12161 | 0.11621 | 0.3320 | 0.3219 | 0.08271 | 0.07935 |
| 6 | 32 | 2 | 0.12659 | 0.12121 | 0.3395 | 0.3295 | 0.08694 | 0.08342 |
| 8 | 64 | 2 | 0.12721 | 0.12181 | 0.3404 | 0.3304 | 0.08745 | 0.08392 |
| 5 | 32 | 2 | 0.12899 | 0.12354 | 0.3430 | 0.3330 | 0.08890 | 0.08533 |

### 3 Performance metrics

To evaluate the deep learning models, we used standard performance metrics appropriate for each task. For the pattern-classification problem, we considered categorical accuracy (CA), mean intersection-over-union (IoU), area under the receiver operating characteristic curve (ROC AUC), and categorical cross-entropy loss (CCE). For the regression problem of estimating effective mortality, we used mean squared error (MSE), mean absolute error (MAE), and mean squared logarithmic error (MSLE).

##### 3.1 Categorical accuracy

Categorical accuracy measures the fraction of correctly classified samples:

$$CA = \frac{\text{number of correctly classified samples}}{\text{total number of samples}} = \frac{1}{n}\sum_{i=1}^{n} \mathbb{I}(\hat{y}_i = y_i), \tag{3.1}$$

where $n$ is the total number of samples, $y_i$ is the true class label of sample $i$, $\hat{y}_i$ is the predicted class label, and $\mathbb{I}$ is the indicator function. A value of 1 indicates perfect classification.

##### 3.2 Mean intersection-over-union (IoU)

For the multi-class classification problem, we computed the mean IoU across classes. For a given class $j$, IoU is defined as

$$IoU_j = \frac{TP_j}{TP_j + FP_j + FN_j}, \tag{3.2}$$

Here, $TP_j$, $FP_j$, and $FN_j$ denote the true positives, false positives, and false negatives for class $j$, respectively. The mean IoU is then obtained by averaging over all classes:

$$IoU = \frac{1}{c}\sum_{j=1}^{c} IoU_j, \tag{3.3}$$

where $c$ is the number of classes. Values closer to 1 indicate better agreement between predicted and true labels.

##### 3.3 Area under the receiver operating characteristic curve (ROC AUC)

The ROC AUC measures the ability of a classifier to rank the true class above alternative classes over the full range of decision thresholds [12]. Because our problem is multiclass, we used a micro-averaged ROC AUC that computes the metric globally across all samples and classes. For details on the implementation, refer to the documentation of the corresponding *scikit-learn* routine [13]. Higher ROC AUC values indicate better discrimination performance.

##### 3.4 Categorical cross-entropy loss (CCE)

Categorical cross-entropy measures the discrepancy between the predicted class probabilities and the true labels:

$$CCE = -\frac{1}{n}\sum_{i=1}^{n}\sum_{j=1}^{c} y_{i,j}\log(p_{i,j}), \tag{3.4}$$

where $n$ is the total number of samples, $c$ is the total number of classes, $y_{i,j}$ is the one-hot encoded true label, and $p_{i,j}$ is the predicted probability that sample $i$ belongs to class $j$. Lower values indicate better classification performance.

##### 3.5 Mean squared error (MSE)

For mortality estimation, the main optimization metric was the mean squared error,

$$MSE = \frac{1}{n}\sum_{i=1}^{n}(w_i - \hat{w}_i)^2, \tag{3.5}$$

where $n$ is the total number of samples, $w_i$ is the true effective mortality of sample $i$, and $\hat{w}_i$ is the predicted value. An MSE of 0 indicates a perfect prediction.

##### 3.6 Mean absolute error (MAE)

The mean absolute error is defined as

$$MAE = \frac{1}{n}\sum_{i=1}^{n} |w_i - \hat{w}_i|, \tag{3.6}$$

and measures the mean absolute difference between the predicted and true values, respectively.

##### 3.7 Mean squared logarithmic error (MSLE)

The mean squared logarithmic error is defined as

$$MSLE = \frac{1}{n}\sum_{i=1}^{n} \left[\log(w_i + 1) - \log(\hat{w}_i + 1)\right]^2, \tag{3.7}$$

and measures the mean-squared difference between the logarithms of the predicted and true values. This metric is only defined when both $w_i$ and $\hat{w}_i$ are non-negative.

#### 4 Empirical dataset compilation

##### 4.1 Details of the empirical dataset

The benthic cartography covers approximately 2,500 km$^2$ of shallow coastal habitats and was produced over roughly two decades through a combination of European and regional monitoring projects, with a major update completed in 2018. It was derived primarily from side-scan sonar, complemented by photo-interpretation of airborne imagery and in situ observations, providing a high-resolution representation of benthic spatial configuration. Habitat classes follow the Standard List of Marine Habitats of Spain (LPHME). The 28 original classes were first aggregated into four broader ecological categories representative of Mediterranean coastal systems—*Posidonia oceanica*, Other green plants, Brown algae & rocks and Sandy bottoms [14, 15]—before being further reduced to a binary seagrass presence-absence map for use in the present analysis.

##### 4.2 Empirical habitat cartography composition

The original seabed cartography contains a total of 28 different classes, which were aggregated into 4 major ecological groups or habitat types (*Posidonia oceanica*, Other green plants, Brown algae & rocks and Sandy bottoms) based on feature similarity and ecological function (Supplementary Table 7). Although these habitat classes are present in the whole Mediterranean Sea, the specific composition of the underlying sub-classes can vary among the different islands (e.g., one particular species might be present on only one island, such as *Zostera noltii*).

**Supplementary Table 7.** Ecological Categories and Subcategories in the ground truth habitat data.

| **Category** | **Subcategory** | **Area (km$^2$)** | **Presence zone** |
|---|---|---|---|
| **Posidonia oceanica** | *Posidonia oceanica* | 538.61 | Mallorca, Menorca, Ibiza, Formentera |
| | Barrier reef of *Posidonia oceanica* | 0.49 | Mallorca, Menorca |
| | *Posidonia oceanica* on stone with sand | 20.33 | Mallorca, Menorca |
| | Mixed *Posidonia oceanica* with dead rhizome | 0.10 | Ibiza |
| | Meadows of *Posidonia oceanica* on dead mat (rhizome) | 5.23 | Mallorca, Menorca |

| | | | |
|---|---|---|---|
| **Other Green Plants** | Algae photophilic on stone with *Posidonia oceanica* | 6.63 | Mallorca, Menorca |
| | *Caulerpa prolifera* | 0.82 | Mallorca, Menorca, Ibiza, Formentera |
| | Meadows of phanerogams and green rhizomatous algae | 4.87 | Mallorca, Menorca |
| | Fine sands with *Cymodocea nodosa* | 1.95 | Mallorca, Menorca, Ibiza, Formentera |
| | *Cymodocea nodosa* | 1.86 | Mallorca, Menorca, Ibiza, Formentera |
| | *Zostera noltii* | 0.01 | Menorca |
| | *Cymodocea nodosa* and *Zostera noltii* | 0.04 | Menorca |
| | Mixed meadows of *Cymodocea nodosa* and *Caulerpa prolifera* | 4.62 | Mallorca, Menorca, Ibiza |
| **Sandy Bottoms** | Coarse sands | 250.19 | Ibiza, Formentera |
| | Soft or sedimentary substrate | 584.71 | Mallorca, Menorca |
| | Mud | 0.03 | Ibiza |
| | Fine sands | 74.36 | Ibiza, Formentera |
| | Muddy detrital bottom | 11.52 | Ibiza |
| | Leptometra phalangium fields in bathyal bottoms of platform edge | 208.85 | Menorca |
| | Bathyal bottoms of platform edge with *Gryphus vitreus* | 91.93 | Menorca |
| | Medium sands | 42.41 | Ibiza, Formentera |
| | Muddy detrital bottoms infralittoral and circalittoral | 589.81 | Mallorca, Menorca, Formentera |
| **Brown Algae and Rocks** | Rocky bottoms with photophilic algae and sands | 36.49 | Mallorca, Menorca |
| | Rocky bottoms dominated by sciafilic and hemisciafilic algae | 21.17 | Mallorca, Menorca, Ibiza, Formentera |
| | Cliffs, walls, and rocky slopes of the deep sea | 13.96 | Menorca |
| | Rocky bottoms with photophilic algae | 14.35 | Ibiza, Formentera |
| | Rocky bottoms with photophilic algae and sands | 15.58 | Mallorca, Menorca |

#### 4.3 Dataset creation

To classify the different patterns present in the Balearic Islands and estimate their associated mortality, the data were divided into patches. These patches were created with the same dimensions as those used to train the neural networks, each of $256 \times 256$ pixels. Because the data resolution is 1 meter per pixel, each of our patches covers an area of approximately 6.5 hectares. Predictions were made for the coasts of the four main islands of the Balearic Islands: Mallorca, Menorca, Ibiza, and Formentera.

#### 4.4 Details about the spatial indicators

Seagrass cover was defined as the fraction of pixels occupied by *P. oceanica* within each image tile, providing a first-order measure of meadow occupancy. The mean hole area was computed for samples classified as meadows with holes, where holes were identified as connected components of non-vegetated pixels fully enclosed within the seagrass matrix; their areas were calculated in square meters and averaged per tile. The mean inter-patch distance was computed for fragmented samples, particularly those in the striped and spotted states. Disconnected seagrass patches were identified as connected components of vegetated pixels, and the minimum distances between all patch pairs were calculated; the mean nearest-neighbor distance was used as an indicator of patch isolation, with higher values corresponding to greater spatial separation among remnant patches and a more advanced degree of fragmentation.

### 5 Model test results

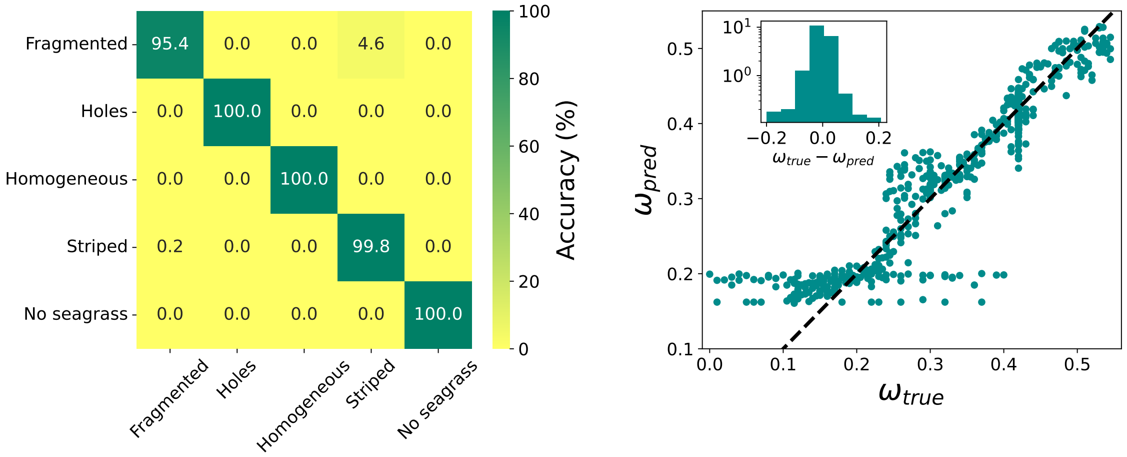

**Supplementary Fig. 11. Performance of the CNN models on the held-out synthetic test set.** (a) Confusion matrix for the pattern-classification model, showing the correspondence between true and predicted meadow states. Misclassifications are rare and occur mainly between neighboring states along the model-defined deterioration sequence. (b) Predicted versus true effective mortality $\omega$ for the regression model. The dashed line indicates perfect agreement, and the inset shows the distribution of prediction errors. Regression performance is high across most of the parameter range, with larger errors at low effective mortality, where binarization removes density information and makes homogeneous states with different values of $\omega$ less distinguishable.

### 6 Pattern and mortality inference

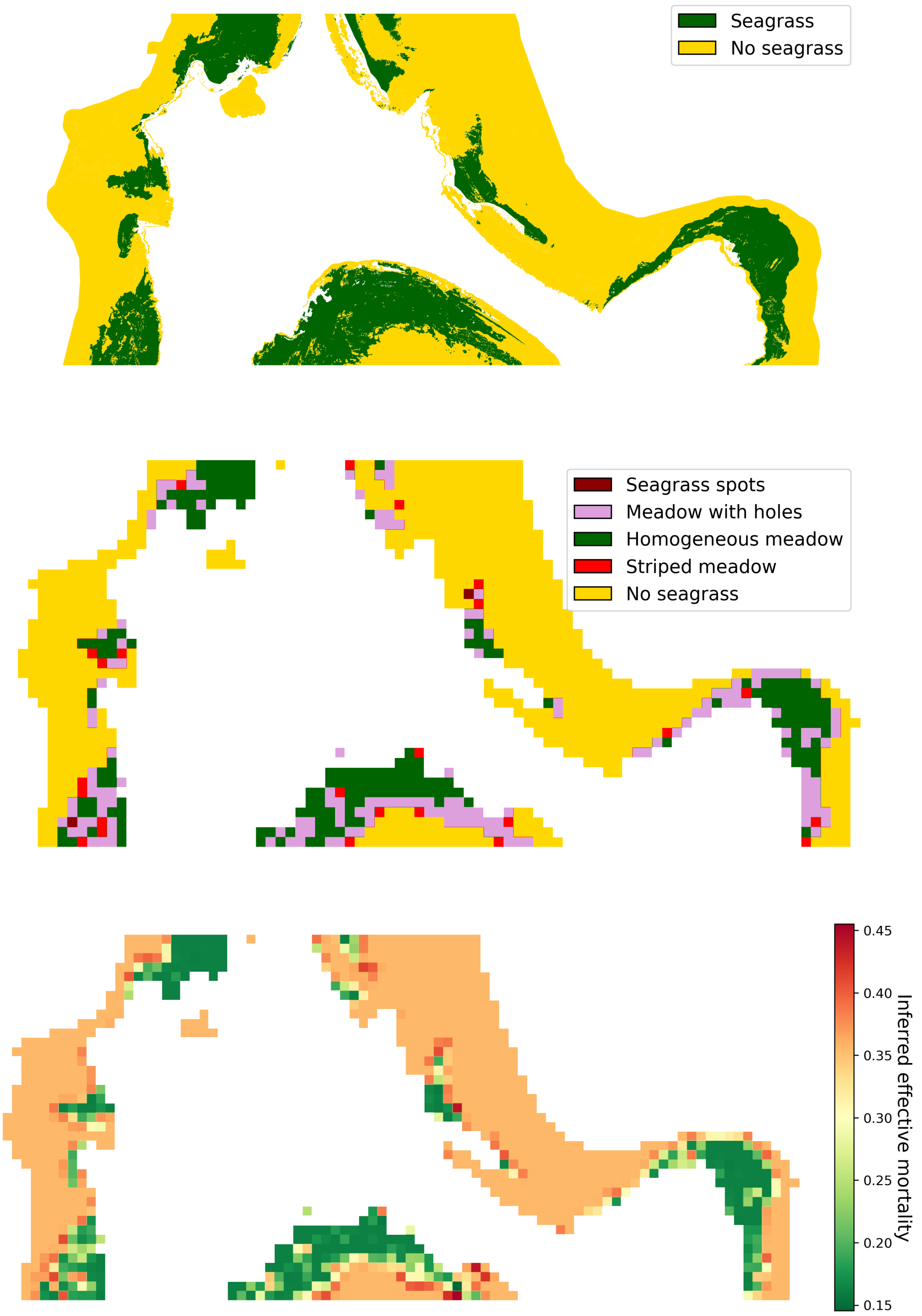

**Supplementary Fig. 12. Example of spatially explicit meadow-state classification and inferred effective mortality in northern Formentera.** Top: Binary empirical habitat cartography showing the distribution of *P. oceanica* and non-seagrass habitats. Middle: Discrete meadow states inferred by the classification model, including homogeneous meadow, meadow with holes, striped meadow, seagrass spots, and no seagrass. Bottom: spatially explicit map of inferred effective mortality obtained using the regression model for the same area. The mortality map extends the discrete classification into a continuous estimate of meadow deterioration, revealing fine-scale spatial variation in condition within and along seagrass patches.

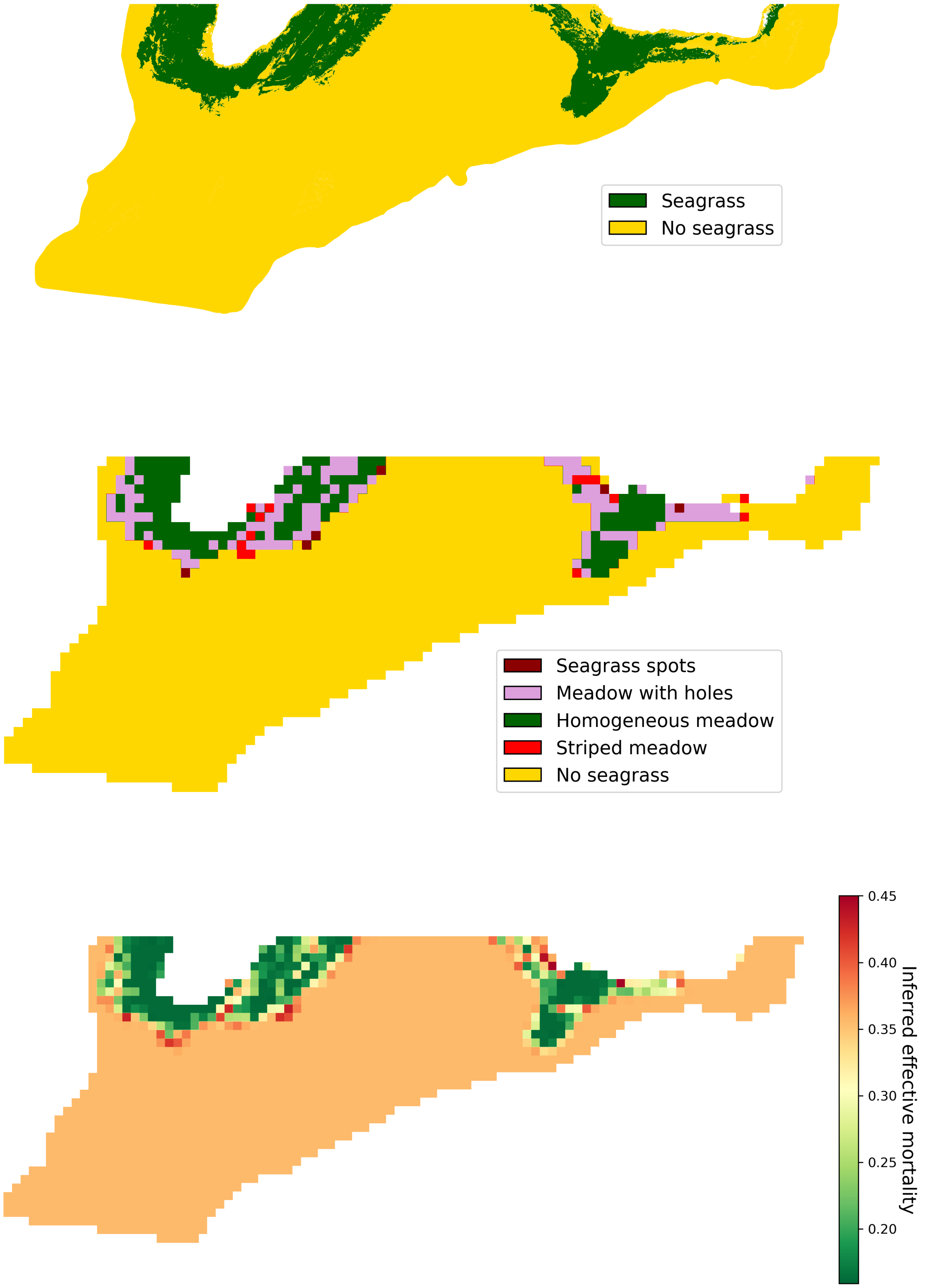

**Supplementary Fig. 13. Additional example of spatially explicit meadow-state classification and inferred effective mortality in southern Formentera.** Top: Binary empirical habitat cartography showing the distribution of *P. oceanica* and non-seagrass habitats. Middle: discrete meadow states inferred using the classification model. Bottom: corresponding map of inferred mortality parameter $\omega$ from the regression model.

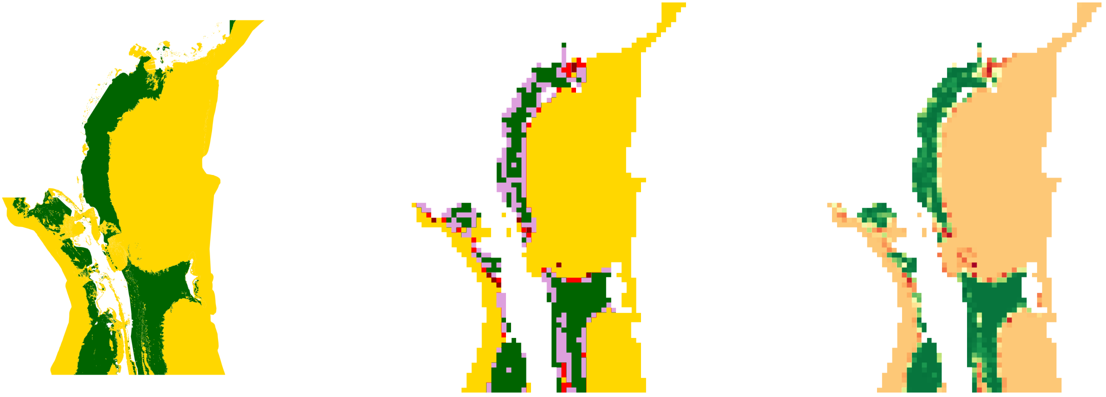

**Supplementary Fig. 14. Additional example of spatially explicit meadow-state classification and inferred effective mortality in southern Ibiza.** Left: Binary empirical habitat cartography. Middle: Discrete meadow states inferred using the classification model. Right: Map of inferred mortality parameter $\omega$ from the regression model. Color codes are the same as in the previous figures.

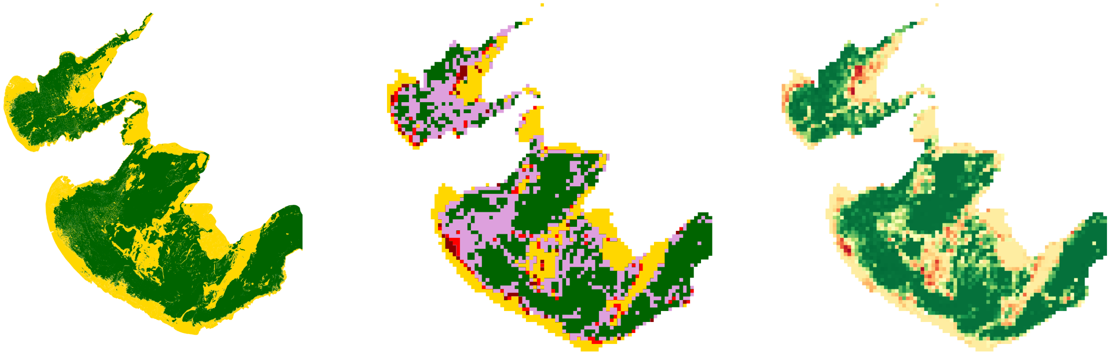

**Supplementary Fig. 15. Additional example of spatially explicit meadow-state classification and inferred effective mortality in northeast Mallorca.** Left: Binary empirical habitat cartography. Middle: Discrete meadow states inferred using classification model. Right: Corresponding map of the inferred effective mortality from the regression model. Color codes are the same as in the previous figures.

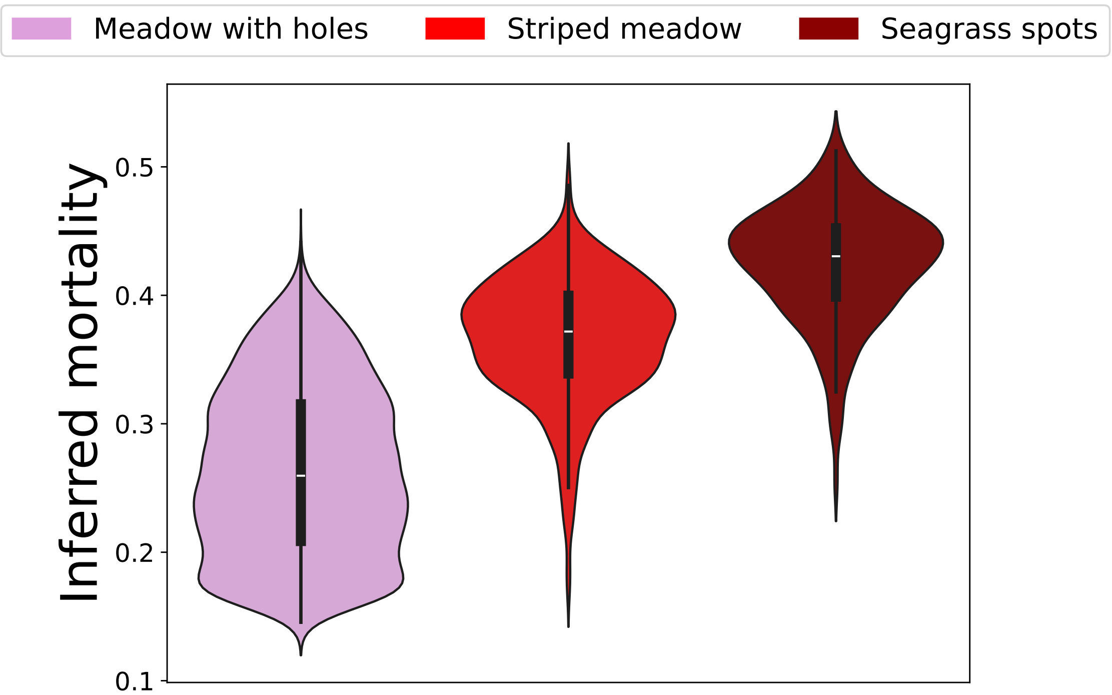

**Supplementary Fig. 16. Distribution of inferred mortality across degraded pattern classes.** Violin plots of inferred mortality for samples classified as meadows with holes, striped meadows, and seagrass spots. The distributions follow the expected order along the deterioration gradient, with lower values in meadows with holes and progressively higher values in striped meadows and seagrass spots. This supports the interpretation that the inferred mortality captures a meaningful latent axis of meadow condition.

### 7 Comparison with null model

To test whether CNN inference depended only on seagrass cover, we generated a randomized counterpart of each empirical binary image by randomly permuting its pixels. This procedure preserved image dimensions and seagrass cover exactly while destroying spatial organization. The randomized images were then processed using the same classification and regression models as the original images. The resulting changes in pattern classification and inferred effective mortality are shown in Fig. 5 of the main text.

Across the 14 study zones, observed and randomized class abundances differed significantly for all five states according to paired Wilcoxon signed-rank tests ($p \leq 0.0039$; Supplementary Table 8). The largest changes occurred for the patterned states. A secondary Mann–Whitney comparison, which does not account for the paired structure of the data, is reported for completeness. Because each original–randomized image pair has exactly the same seagrass cover, these results show that both classification and inference of the effective mortality parameter depend on spatial organization beyond vegetation occupancy alone.

**Supplementary Table 8.** Comparison of observed and cover-preserving randomized classifications across the 14 study zones. The Mann–Whitney $U$ test treats observed and randomized counts as independent samples, whereas the Wilcoxon signed-rank test accounts for their paired structure within zones and is therefore the primary analysis. Wilcoxon $p$-values were adjusted for the five pattern-class comparisons using the Holm procedure.

| Pattern state | Mann–Whitney $U$ | $p_{\mathrm{MW}}$ | Wilcoxon $W$ | $p_{\mathrm{W}}$ | Holm-adjusted $p$ |
|---|---|---|---|---|---|
| Seagrass spots | 196.00 | $< 0.0001$ | 0.00 | 0.0009674 | 0.00387 |
| Meadow with holes | 196.00 | $< 0.0001$ | 0.00 | 0.0009787 | 0.00387 |
| Homogeneous meadow | 74.00 | 0.2802 | 0.00 | 0.0009787 | 0.00387 |
| Striped meadow | 196.00 | $< 0.0001$ | 0.00 | 0.0009759 | 0.00387 |
| No seagrass | 80.00 | 0.4213 | 0.00 | 0.0001221 | 0.0006104 |

### 8 State-specific uncertainty of inferred mortality

To quantify uncertainty in the inferred mortality parameter, we used the synthetic test set, which was not used during model training or validation and for which the true value of $\omega$ was known. For each test image i, we calculated the residual

$$e_i = \omega_i - \hat{\omega}_i \tag{8.1}$$

Residual distributions were analysed separately for each spatial pattern state because the information available for estimating $\omega$ differs among pattern morphologies. For each state, we calculated the mean residual bias, mean absolute error, root-mean-square error, median absolute error, and the 95th percentile of the absolute residual distribution. We also calculated the 2.5th and 97.5th percentiles of the signed residuals to characterize possible asymmetry in prediction errors. We used these residual distributions to estimate the expected prediction error associated with empirical values of $\hat{\omega}$. The resulting intervals quantify uncertainty in the CNN inverse mapping conditional on the mechanistic model.

Prediction accuracy differed substantially among spatial pattern states (Supplementary Fig. 17 and Supplementary Table 9). The mortality parameter was inferred most precisely for striped meadows, which showed a median absolute error of 0.0094, an RMSE of 0.0166, and a 95th percentile absolute error of 0.0345. Errors were moderately larger for seagrass spots and meadows with holes. Their median absolute errors were 0.0193 and 0.0218, respectively, while their 95th percentile absolute errors were 0.0722 and 0.0659.

Homogeneous meadows showed substantially greater uncertainty, with a median absolute error of 0.1026, an RMSE of 0.1176, and a 95th percentile absolute error of 0.1933. This lower identifiability is consistent with the behaviour observed in the regression results: after binarization, homogeneous landscapes contain little spatial structure with which to distinguish among nearby mortality values, a limitation that is independent of the number of samples per class.

Mean prediction bias was small for all pattern states. The largest bias occurred for meadows with holes ($-0.0206$), followed by seagrass spots ($-0.0108$), indicating a modest tendency to overestimate $\omega$ in these states. Bias was close to zero for striped and homogeneous meadows. Overall, the results indicate that mortality inference is precise within the patterned regime, particularly for striped configurations, whereas estimates for homogeneous meadows should be interpreted with greater caution.

**Supplementary Table 9.** State-specific prediction errors of the mortality regressor evaluated on the held-out synthetic test set. Residuals were defined as $e = \omega_{\mathrm{true}} - \hat{\omega}_{\mathrm{pred}}$. The last two columns show the 2.5th and 97.5th percentiles of the signed residual distribution.

| Pattern state | $n$ | Bias | MAE | Median AE | RMSE | $q_{0.95}(\lvert e\rvert)$ | $q_{0.025}(e)$ | $q_{0.975}(e)$ |
|---|---|---|---|---|---|---|---|---|
| Striped meadow | 508 | 0.0019 | 0.0122 | 0.0094 | 0.0166 | 0.0345 | -0.0267 | 0.0436 |
| Seagrass spots | 211 | -0.0108 | 0.0251 | 0.0193 | 0.0330 | 0.0722 | -0.0806 | 0.0447 |
| Meadow with holes | 102 | -0.0206 | 0.0283 | 0.0218 | 0.0353 | 0.0659 | -0.0761 | 0.0247 |
| Homogeneous meadow | 57 | 0.0051 | 0.1052 | 0.1026 | 0.1176 | 0.1933 | -0.1774 | 0.1977 |

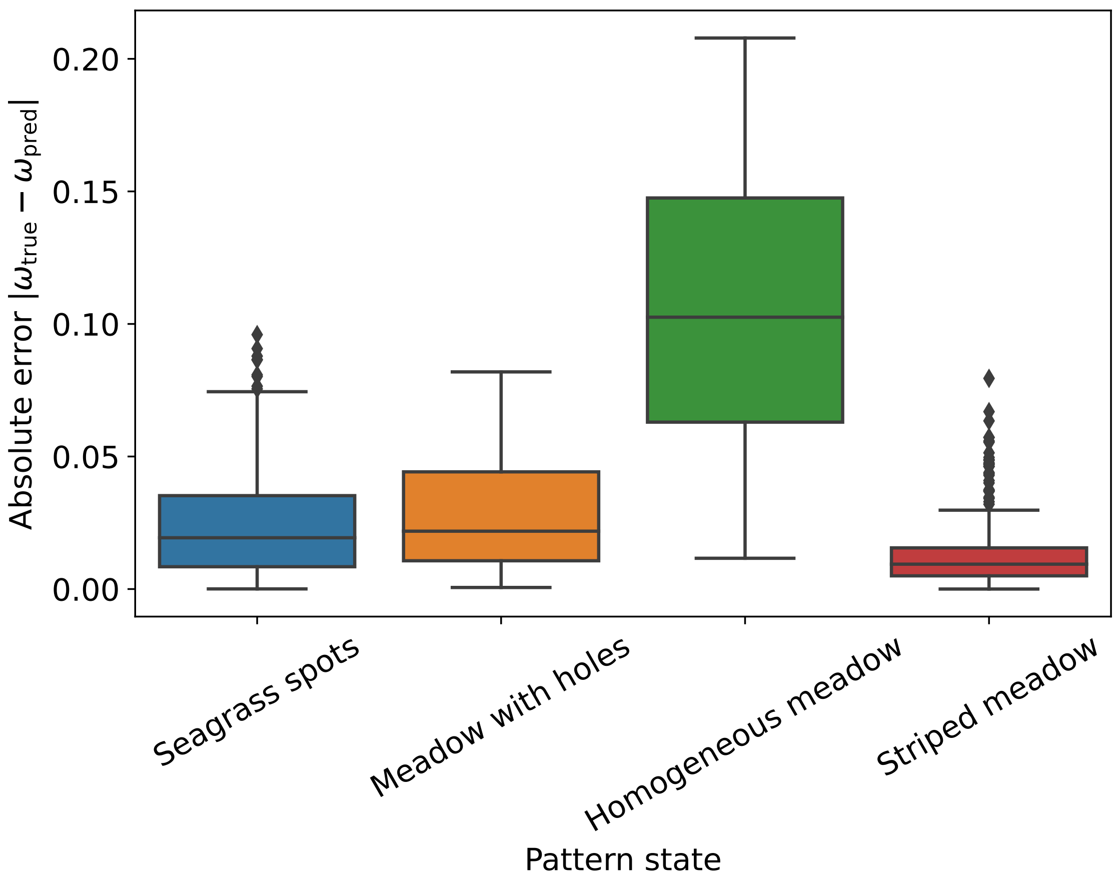

**Supplementary Fig. 17. State-specific prediction error of the mortality regressor.** Boxplots show the distribution of the absolute error $|\omega_{\mathrm{true}} - \hat{\omega}_{\mathrm{pred}}|$ in the held-out synthetic test set, stratified by spatial pattern state. Prediction errors were lowest for striped meadows and highest for homogeneous meadows, for which binarization removes most of the density information available to distinguish among values of $\omega$.
